# Mechanistic basis of temperature-adaptation in microtubule dynamics across frog species

**DOI:** 10.1101/2024.07.29.605571

**Authors:** Luca Troman, Ella de Gaulejac, Abin Biswas, Jennifer Stiens, Benno Kuropka, Carolyn Moores, Simone Reber

**Author notes:** Contributed equally (Luca Troman, Ella de Gaulejac).

## Abstract

Cellular processes are remarkably effective across diverse temperature ranges, even with highly conserved proteins. In the context of the microtubule cytoskeleton, which is critically involved in a wide range of cellular activities, this is particularly striking as tubulin is one of the most conserved proteins while microtubule dynamic instability is highly temperature sensitive. We thus lack a mechanistic framework that links functional adaptability with environmental pressures. Here, we leverage the diversity of natural tubulin variants from three closely related frog species that live at different temperatures: we combine *in vitro* reconstitution assays, quantitative biochemistry, and cryogenic electron microscopy to show how a small number of primary sequence changes influences the energy landscape of tubulin interactions and thereby mediates cold-adaptation and microtubule stability. This study thus broadens our conceptual framework for understanding microtubule dynamics and provides insights into how conserved cellular processes are tailored to different ecological niches.

## INTRODUCTION

The body temperature of ectotherms is largely determined by their habitat temperature. As they cannot actively regulate their body temperature, ectotherms are considered to be particularly vulnerable to environmental temperature fluctuations. On a cellular level, temperature is an important environmental variable impacting all forms of biological rates and functions. A cellular structure that is known to be exquisitely cold sensitive is the microtubule cytoskeleton, which quickly disappear on incubation at low temperatures.^1–5^ Microtubules are dynamic polymers of αβ-tubulin with essential cellular functions including cell division, motility, and signalling. How did microtubules adapt to temperature to ensure function in species that live across a considerable temperature range? Dissecting the adaptations of microtubules to diverse temperature ranges across species thus addresses a fundamental challenge in understanding how proteins involved in conserved cellular processes are tailored to different ecological niches.

The microtubule’s cold sensitivity is a consequence of how microtubules assemble: they grow longitudinally by the addition of GTP-associated tubulin into protofilaments (pf), a process known to be largely driven by the entropically favourable displacement of structured water at the subunit interfaces.^5–7^ Protofilaments then link via lateral connections to form a cylindrical, polar microtubule. The bound GTP is hydrolysed to GDP, leading to conformational changes in tubulin that render the microtubule lattice less stable and eventually result in the microtubule depolymerizing. This transition from growth to shrinking is termed catastrophe while the transition from shrinking to growth is termed rescue. This stochastic switching between growing and shrinking states is known as “dynamic instability”,^8^ a non-equilibrium behaviour with nontrivial temperature dependencies. Detailed mechanistic insights about dynamic instability have come from *in vitro* reconstitution of microtubules with purified components. While many advances stem from reconstituted microtubules from endotherm tubulin purified mainly from porcine or bovine brains, the majority of animals on earth are ectotherms.

One molecular strategy that cells can use to cope with disruptive effects of temperature changes on microtubule dynamics involves microtubule-associated proteins (MAPs). In mammals, for example, the microtubule-associated protein MAP6 has been shown to act as temperature sensor and protect microtubules against cold-induced depolymerization.^9^ In the absence of such MAPs mammalian microtubules depolymerize below 20°C. Some cold-adapted species such as Antarctic fishes and glacier ice worms, however, assemble microtubules even below 10°C.^10–13^ In these cryophilic ectotherms, tubulin – despite being highly conserved across eukaryotes^14^ – shows species-specific amino acid substitutions mostly in the β-tubulin gene. Consistently, mutation screens in bakers yeast have identified single amino acid changes that turn microtubules cold stable^15^ or cold sensitive^16^. This suggests that a small set of rare residue substitutions confers the ability to polymerize microtubules at low physiological temperatures. Indeed, the incorporation of a single additional tubulin isotype can confer cold-tolerance to cytoplasmic microtubules in human cells^12^ and flies^17^. As these studies have been carried out on very divergent organisms, it remains difficult to pinpoint how sequence differences manifest in microtubule growth at low temperatures. We thus miss a unifying picture and systematic understanding of microtubule cold-adaptation.

To gain such a systematic understanding of how a small number of primary sequence changes can mediate cold-adaptation of microtubule dynamics, we studied microtubules from three *Xenopus* species that have evolved to live at significantly different temperatures ranging from 18-27°C (Figure 1A). We reconstituted microtubule growth *in vitro* and found that *Xenopus* tubulin orthologues evolved to exhibit comparable polymerization velocities but significantly different catastrophe frequencies and rescue probabilities at divergent thermal niches. Using these “natural tubulin mutants”, we determined the structures of microtubules across all three species to overall resolutions better than 4 Å. In our data, *Xenopus* tubulin polymerizes to form mostly non-canonical 14- or 15-protofilament microtubules. We found the GDP-tubulin lattices to be very similar across the three species with small differences in the β-tubulin lateral interactions. We propose that more flexible lateral contacts in cold-adapted frogs have an additive effect that stabilises microtubules. At the dynamic microtubule plus-end, we found the rates of GTP hydrolysis to be identical. The free energy of tubulin incorporation, however, scaled inversely with ambient temperature. This means that the cold-adapted *X. laevis* tubulin has the highest free energy in solution and thus readily incorporates into the growing microtubule plus-end explaining fast microtubule growth velocity and low critical concentration. In summary, these results elucidate the structure-dynamics-function relationship associated with the strategies of thermal adaptation of microtubules.

**Figure 1.**
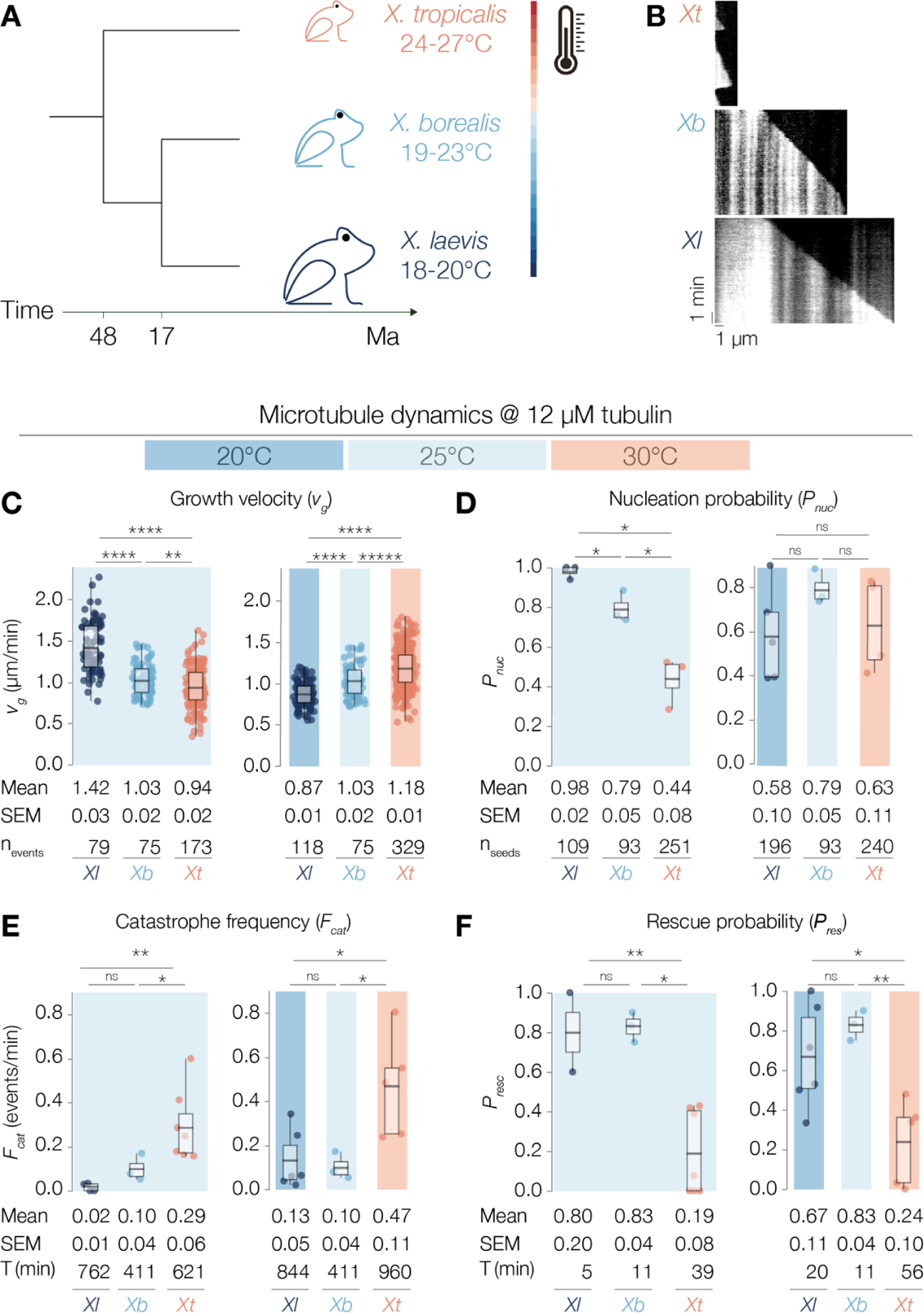
Tubulin orthologues in closely related *Xenopus* species evolved to exhibit comparable polymerization velocities but significantly different catastrophe frequencies and rescue probabilities at divergent thermal niches. (**A**) Schematic depicting *Xenopus* phylogeny^73^ and optimal breeding temperatures. (**B**) Representative TIRF kymographs showing dynamic *Xl*, *Xb*, and *Xt* microtubules at 12 µM tubulin and 25°C. (**C-F**) Parameters of dynamic instability. All values were obtained from measurements of microtubules pooled over at least 3 independent experiments, all p-values were calculated with the Mann-Whitney test, g corresponds to Hedge’s g. Data points correspond to individual measurements (C, D) or experimental replicates (E, F). All error values are SEM. Plot boxes range from 25th to 75th percentile, whiskers span the range, horizontal line shows median value. (**C**) At 25°C, *Xl* microtubules grow at 1.42 ± 0.03 µm/min, *Xb* at 1.03 ± 0.02 µm/min and *Xt* at 0.94 ± 0.02 µm/min, with p-value(*Xl,Xb*)<0.0001, g(*Xl,Xb*)=1.53; p-value(*Xb,Xt*)=0.01, g(*Xb,Xt*)=0.38; p-value(*Xl,Xt*)<0.001, g(*Xl,Xt*)=1.84. Right graph: At 20°C, *Xl* microtubules grow at 0.87 ± 0.01 µm/min, 25°C, *Xb* at 1.03 ± 0.02 µm/min and 30°C, *Xt* at 1.18 ± 0.01 µm/min, with p-value(*Xl,Xb*)<0.0001, g(*Xl,Xb*)=0.94; p-value(*Xb,Xt*)<0.0001, g(*Xb,Xt*)=0.67; p-value(*Xl,Xt*)<0.0001, g(*Xl,Xt*)=1.40. (**D**) Nucleation probability is the percentage of seeds that produced at least one microtubule within 10 min of acquisition^23^ on the total number of seeds in the field of view. At 25°C, *Xl* microtubules have a nucleation probability of 0.98 ± 0.02, *Xb* of 0.79 ± 0.05, *Xt* of 0.44 ± 0.08, with p-value(*Xl,Xb*)=0.04; p-value(*Xb,Xt*)=0.02; p-value(*Xl,Xt*)=0.01. Right graph: At 20°C, *Xl* microtubules have a nucleation probability of 0.58 ± 0.10, 25°C *Xb* microtubules of 0.79 ± 0.05, 30°C *Xt* microtubules of 0.63 ± 0.11, with p-value(*Xl,Xb*)=0.10; p-value(*Xb,Xt*)=0.24; p-value(*Xl,Xt*)=0.74. n_seeds_ is the total number of seeds used for the analysis. (**E**) Catastrophe frequencies are reported as the total number of catastrophes over the total time spent growing. At 25°C, *Xl* microtubules catastrophe at a frequency of 0.02 ± 0.01 events/min, *Xb* microtubules 0.10 ± 0.04 events/min, *Xt* microtubules 0.29 ± 0.06 events/min, with p-value(*Xl,Xb*)=0.14; p-value(*Xb,Xt*)=0.03; p-value(*Xl,Xt*)<0.01. Right graph: At 20°C, *Xl* microtubules catastrophe at a frequency of 0.13 ± 0.05 events/min, 25°C *Xb* microtubules 0.10 ± 0.04 events/min, 30°C *Xt* microtubules 0.47 ± 0.11 events/min, with p-value(*Xl,Xb*)=0.69; p-value(*Xb,Xt*)=0.02; p-value(*Xl,Xt*)=0.03. T(min) is the total time microtubules spent growing. (**F**) Rescue probability is the ratio of the total number of rescued catastrophes over the total number count of all catastrophes. At 25°C, *Xl* microtubules have a rescue probability of 0.80 ± 0.20 (2 of 4 replicates without catastrophe), *Xb* microtubules of 0.83 ± 0.04, *Xt* microtubules of 0.19 ± 0.08, with p-value(*Xl,Xb*)=0.91; p-value(*Xb,Xt*)<0.001; p-value(*Xl,Xt*)=0.16. Right graph: At 20°C, *Xl* microtubules have a rescue probability of 0.67 ± 11, at 25°C *Xb* microtubules of 0.83 ± 0.04, 30°C *Xt* microtubules of 0.24 ± 0.10, with p-value(*Xl,Xb*)=0.20; p-value(*Xb,Xt*)<0.01; p-value(*Xl,Xt*)=0.02. T(min) is the total microtubules spent depolymerizing. See also Figures S1.

## RESULTS

### Microtubules show temperature-adaptive differences in closely related *Xenopus* species

Microtubule dynamics are highly sensitive to changes in temperature. To compare how microtubules of closely related ectothermic *Xenopus* species (Figure 1A) adapt to temperature, we purified tubulin^18–20^ from extracts prepared from unfertilized *X. laevis*, *borealis*, and *tropicalis* eggs (Figure S1A). Using a total internal reflection microscopy (TIRFM)-based *in vitro* assay^21^ and kymograph analysis (Figure 1B),^22^ we measured intrinsic microtubule plus-end dynamics from GMPCPP-stabilized microtubule seeds, free of MAPs and motors. At 25°C and 12 µM tubulin, *X. laevis* microtubules showed an increased polymerization velocity (Figure 1C, v_g_Xl_ = 1.42 ± 0.03 µm/min, arithmetic mean ± SEM), a higher stochastic nucleation probability (Figure 1D), defined as the probability of a GMPCPP-seed producing at least one microtubule in t = 10 min, P_nuc_Xl_ = 0.98 ± 0.02,^23^ a lower catastrophe frequency (Figure 1E, F_cat_Xl_ = 0.02 ± 0.01 events/min), and rescued with a greater probability (Figure 1F, P_res_Xl_ = 0.80 ± 0.20) when compared to *X. tropicalis* microtubules (v_g_Xt_ at 0.94 ± 0.02 µm/min, P_nuc_Xt_ = 0.44 ± 0.08, F_cat_Xt_ 0.29 ± 0.06 events/min, P_res_Xt_ = 0.19 ± 0.08) consistent with previous reports.^19^ For polymerization velocity, nucleation probability, and catastrophe frequency, *X. borealis* microtubules showed an intermediate phenotype (v_g_Xb_ at 1.03 ± 0.02 µm/min, P_nuc_Xb_ = 0.79 ± 0.05, F_cat_Xb_ = 0.10 ± 0.04 events/min). Rescue probabilities for *X. borealis* microtubules (P_res_Xb_ = 0.83 ± 0.04), however, were similar to those of *X. laevis*. As the three *Xenopus* species live at different ambient temperatures, we next measured microtubule dynamics at species-specific temperatures (Figure C-F, second panel) and found the polymerization velocity and nucleation probability to become more similar. Catastrophe frequencies and rescue probabilities remained significantly different, in particular between *X. laevis* and *X. tropicalis*. Depolymerization velocity was comparable for all three microtubule species at 25°C (Figure S1B, v_s_Xl_ = 14.04 ± 1.85 µm/min, v_s_Xb_ = 16.46 ± 1.94 µm/min, v_s_Xt_ = 17.28 ± 0.74 µm/min) and increased with increasing temperatures as expected for a thermodynamically-driven process. Collectively, these data suggest that tubulin orthologues evolved to exhibit comparable polymerization velocities but significantly different catastrophe frequencies and rescue probabilities at divergent thermal niches.

### Cold-adapted species combine a fast microtubule polymerization rate with low catastrophe and high rescue frequencies

To systematically study the temperature-adaptive differences in microtubule dynamics, we used a custom-build sapphire stage, which allowed us to control and closely monitor the temperature throughout all TIRF assays (Figure 2A, S2A, M&M). Using this setup, we measured microtubule polymerization velocity at increasing tubulin concentrations for all three *Xenopus* tubulins. As long described,^8^ polymerization velocity increased linearly with tubulin concentration for all three microtubule species (Figure 2B-D). We then systematically determined this relationship at different temperatures. Overall, we observed that *X. laevis* microtubules robustly grew at 15°C and *X. borealis* at 18°C, while *X. tropicalis* microtubules only grew above 25°C (Figure 2E-G, S2C). For all tubulins at all temperatures, polymerization velocity increased linearly with tubulin concentration. At a given temperature, however, the apparent on rate (k_on_) was consistently highest for *X. laevis* and lowest for *X. tropicalis* microtubules (Figure S2B). Thus, microtubule growth from tubulin of three different *Xenopus* species showed significantly different concentration-dependent on-rates.

**Figure 2.**
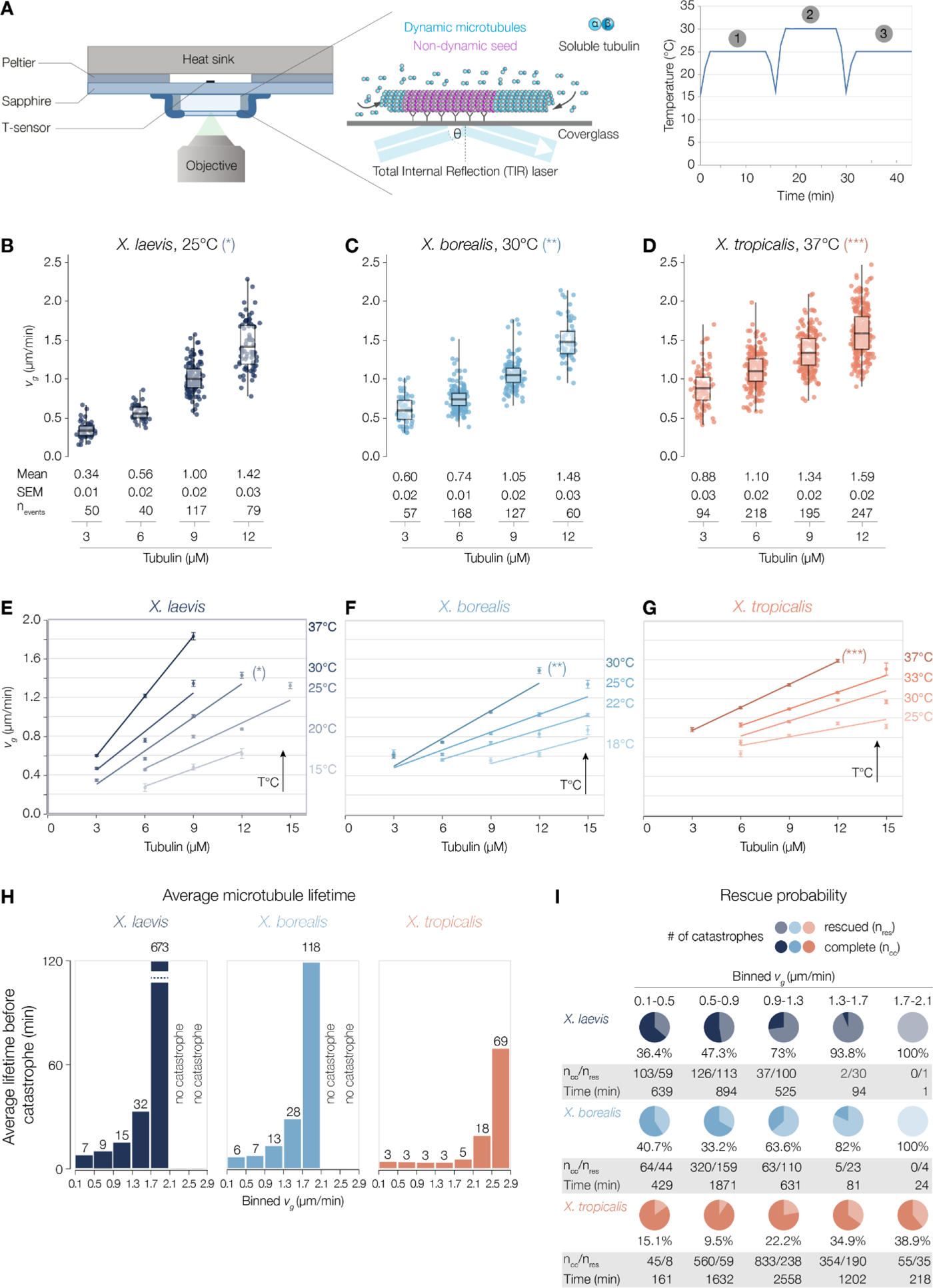
Influence of temperature on *Xenopus* microtubule dynamics. (**A**) Schematic of the experimental setup. (**B-D**) Microtubule growth velocity as a function of tubulin concentration for a single temperature per species. Each datapoint represents a single growth event. Measurements were pooled from at least three independent replicates. Measurements were repeated over a range of temperatures to generate the graphs in (E-G). (**E-G**) Microtubule growth velocity as a function of tubulin concentration for (E) *X. laevis*, (F) *X. borealis* and (G) *X. tropicalis*. Each point is the average growth velocity of measurements pooled over at least 3 independent replicates. Error bars represent SEM. For each temperature, a weighted linear regression was fitted to the data, with weights being the inverse of the variance. Slope of the regression are displayed in Figure S2B. Regression lines marked with (*) in (E), (**) in (F) and (***) in (G) correspond to the data shown in (B), (C) and (D), respectively. (**H**) Average microtubule lifetime binned by growth velocity. Lifetime is defined as the sum of duration of all growth events obtained by pooling measurements from all experiments. For *Xl* and *Xb* no catastrophes were recorded above 2.1 µm/min (time imaged: *Xl* 2.1-2.5 µm/min: 124 min. *Xl* 2.5-2.9 µm/min: 27 min, *Xb* 2.1-2.5 µm/min: 146 min, *Xb* 2.5-2.9 µm/min: 184 min). (**I**) Percentage of rescued versus complete catastrophes for each binned growth velocity. n_cc_ is the count of complete catastrophes, n_res_ of rescues. Time is the sum of the duration of growing events that lead to catastrophe in min. See also Figures S2 and S3.

Because catastrophes are so rare for *X. laevis* and *X. borealis* microtubules (Figure 1E), we – instead of calculating classic catastrophe frequencies (1/s) – measured the microtubule lifetime defined as the average time between two catastrophes binned by microtubule polymerization velocity (Figure 2H). In general, the average microtubule lifetime increased with increasing growth velocities for all three *Xenopus* species. Yet, the average microtubule lifetime at a given growth velocity was significantly higher for *X. laevis* and *X. borealis* when compared to *X. tropicalis*. For example, at a polymerization velocity of 1.3-1.7 µm min^-1^, a *X. laevis* microtubule grew on average 32 minutes before undergoing a catastrophe, a *X. borealis* microtubule 28 minutes but a *X. tropicalis* microtubule only five minutes. Remarkably, at growth velocities above 2.1 µm min^-1^, *X. laevis* and *borealis* microtubules did not catastrophe at all. Thus, while faster growth correlated with longer lifetimes for all three microtubule species (consistent with previous studies^24–26)^, the average lifetime at a given growth velocity significantly differed with *X. laevis* microtubules being the longest-living and *X. tropicalis* the shortest.

Last, to determine the probability of a microtubule being rescued, we determined the number of complete catastrophes (n_cc_) and rescued catastrophes (n_res_) binned by microtubule polymerization velocity. For all three species, the percentage of rescued catastrophes increased with polymerization velocity (Figure 2I). However, the probability of a shrinking microtubule being rescued was significantly higher for *X. laevis* and *X. borealis* when compared to *X. tropicalis*. Similar to catastrophe frequencies, adjusting the temperature close to the respective ambient temperature did not change the significant inter-species differences in rescue probability (Figure 1F). In summary, we show that microtubule growth velocity increases with temperature and tubulin concentration, yet with species-specific on-rates. A higher microtubule growth velocity leads to longer microtubule lifetimes and an increase in rescue probability. Taken together, under identical conditions cold-adapted species combine a fast microtubule polymerization rate with low catastrophe and high rescue frequencies while warm-adapted species show slower polymerization rates with increased catastrophe and decreased rescue frequencies. Next, we set out to understand if these significant differences in dynamic instability can be explained by variations in microtubule architecture.

### Variations in microtubule architecture are not sufficient to explain differences in microtubule dynamics

Microtubules grow by the addition of individual tubulin αβ-heterodimers, aligned longitudinally within protofilaments and linked via lateral connections between protofilaments to form a cylindrical, polar microtubule. In the classic textbook view, microtubules are composed of 13 such protofilaments which, for geometrical reasons^27,28^ form a so-called ‘seam’ (Figure 3A). Although this canonical structure predominates in numerous cellular contexts,^29^ microtubules with divergent architectures have also been described.^30–36^ To explore whether such structural variations exist between our *Xenopus* species, and whether they could contribute to species-specific differences in microtubule dynamics, we prepared samples of polymerized microtubules for investigation using cryo-EM. Importantly, *Xenopus* tubulins have a low critical concentration for microtubule nucleation (Figure 1D), which allowed us to grow the microtubules in the absence of microtubule-stabilising agents, which are known to influence polymer architecture.^37^ Following data acquisition (Figure 3B), we determined the architectural distributions using width measurements of 2D-classified particles and corroborated these with 3D classification (Figure 3C, D, S3A-C, M&M). Surprisingly, each species displayed considerable architectural diversity with the majority of microtubules built from either 14 or 15 protofilaments (Figure 3D). To support these findings, we conducted lattice analysis of individual microtubules using TubuleJ,^38^ which confirmed the presence of both 14-3 and 15-3 microtubules within the sample (Figure 3E). An increase of a single protofilament from 14 to 15 means that 1/14^th^ or 7.14% more dimers need to be incorporated into the lattice per μm of growth, theoretically resulting in a similar change in growth velocity. As our kinetic data showed that growth velocities differed by greater magnitudes, we concluded that the differences in protofilament number were not sufficient to explain the observed growth velocities.

**Figure 3.**
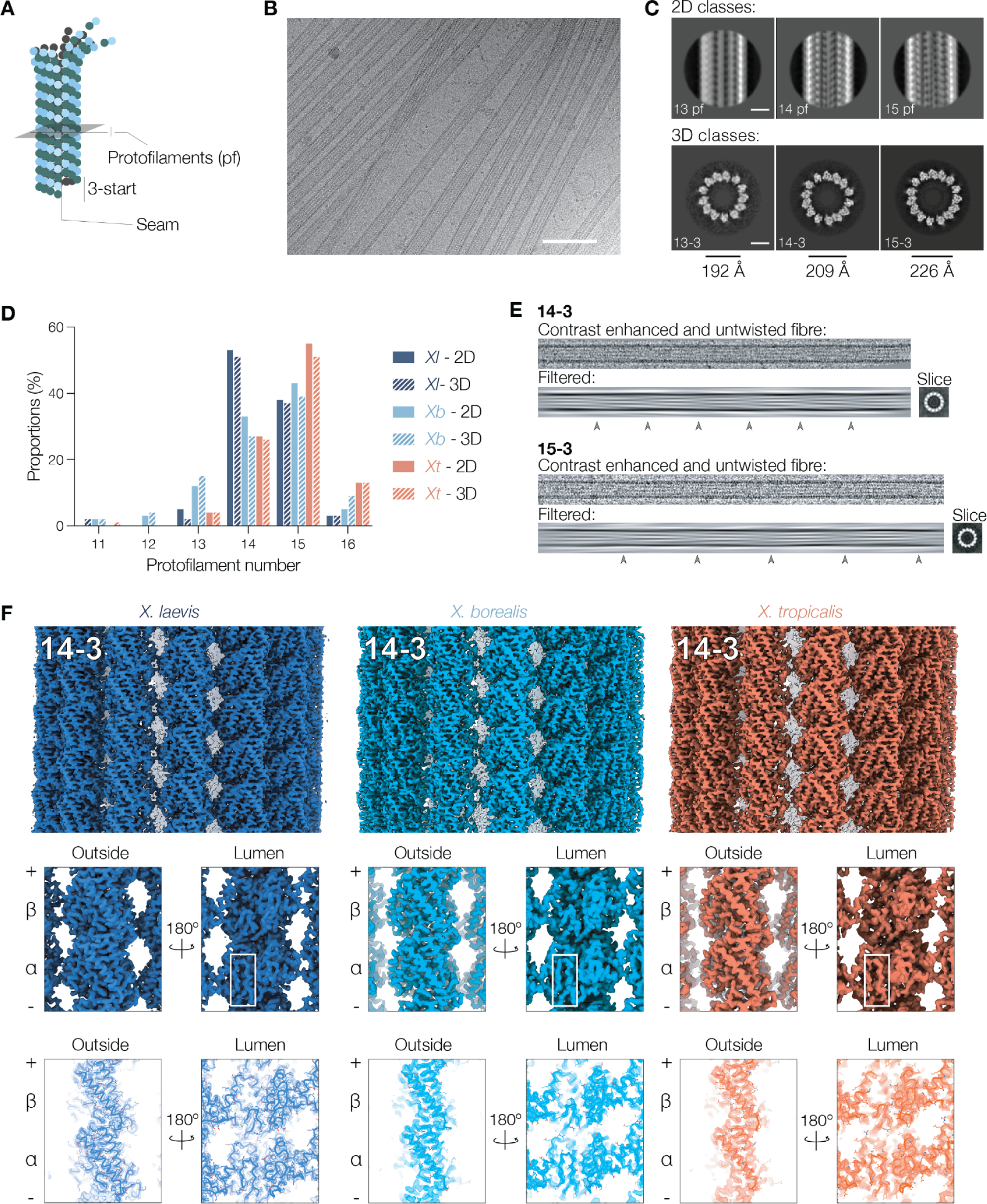
Cryo-EM analysis of microtubules from closely related *Xenopus* species. (**A**) Cartoon diagram of a microtubule with α-tubulin in green and β-tubulin in light blue. (**B**) A representative cryo-EM micrograph of *X. laevis* microtubules. Scale bar = 1000 Å. (**C**) 2D (top) and 3D (bottom) class averages representative of 13, 14, and 15 protofilament microtubules with approximate width measurements shown below. White scale bars = 100 Å. (**D**) Distribution of microtubule protofilament number for all three species determined by 2D and 3D classification. (**E**) Processing of individual microtubules from 2D views using TubuleJ (M&M).^38^ The filtered views (bottom) reveal moiré interference patterns from the protofilaments within the helix (grey arrows). The length and pattern of the repeats indicate underlying protofilament architecture and allows formation of a low resolution 3D reconstruction (transverse cross-sections on the right). (**F**) EM reconstructions of the 14 protofilament 3-start microtubules from each species showing full lattice (top – Figure S4E), single dimers opposite the seam (middle – Figure S4F) and resulting models (bottom – Figure S5). For the lumenal view the appearance of the S9-S10 loop density in α-tubulin opposite the seam (white box) is indicative of particle alignment accuracy, and in particular of seam alignment. Resolutions for all architectures fall between 3.0 and 3.6 Å. See also Figures S4, S5, Table S1.

Previous studies have suggested that fully helical microtubules – lacking the seam – have a higher level of stability.^39–41^ Whilst *X. laevis*, forming the most stable microtubules, had the highest propensity to form helical microtubules (approx. 30% of all microtubules), the equally stable *X. borealis* had the lowest (approx. 15%), and the least stable, *X. tropicalis* microtubules still had approx. 20% fully helical microtubules (Figure S3D). We concluded that in the case of *Xenopus* microtubules, helical symmetry was unlikely to explain the low catastrophe rates observed in the cold-adapted microtubule species. Overall, differences in the intrinsic tendency of tubulins to form microtubules with different protofilament numbers and helical properties were not sufficient to explain the differences in growth velocity and catastrophe frequency between the three *Xenopus* species.

Next, we adapted previously established microtubule processing pipelines (Figure S4, M&M)^42^ to determine the structures of pseudo-helical (3-start) 14- and 15-protofilament microtubules across all three species to 3.0 – 3.6 Å resolutions (Figure 3F). In addition, we solved the structure of the fully helical (4-start) 15-protofilament microtubules in *X. laevis* (Figure S4). Models were built (Figure S5 and below) into each reconstruction allowing us to next explore structural differences between the species on a residue-by-residue basis.

### Subtle structural and allosteric variations must determine microtubule dynamics

Despite its high conservation, tubulin is a genetically and biochemically complex protein family. Thus, to map the correct isoform sequences to the structures, we checked for differential posttranslational modifications and isoform composition in the purified tubulins (Figure 4A). Frog tubulins carried almost no post-translational modifications except for *X. laevis* and *X. borealis* tubulin being phosphorylated (Figure 4B). However, phospho-enrichment coupled with mass spectrometry identified only two very weak phosphosites (Figure S6D). Although the identification of individual tubulin isoforms from *Xenopus* species is challenging (M&M), whole-protein mass spectrometry indicated that the purified tubulins contained three major α-tubulin isoforms (making up more than 80%) and a single β-tubulin (Figure 4C, S6A-C). Sequence alignment revealed a 99% sequence identity within the β-tubulins across the three species and >95% sequence identity within the dominant α-tubulin (mapped onto our experimentally determined structures in Figure 4D, E). Neither of the two amino acid differences between the three species in β-tubulin are directly involved in either nucleotide binding or at sites of tubulin-tubulin lattice interaction (Figure 4D). Although the presence of multiple α-tubulin isoforms within the sample makes interpretation more challenging, mapping of the TUBAL3 sequence in the three species onto their cognate structures showed that amino acid differences were scattered throughout the three-dimensional fold of each subunit rather than concentrated at a specific site. Thus, at the dimer level, all three *Xenopus* tubulins are highly similar. Nevertheless, these few amino acid substitutions must be the origin of the diverse dynamic behaviours, most likely through subtle and allosteric mechanisms. To investigate whether conformational differences within the microtubule lattice offer more insights, we next compared lattice arrays that included longitudinal and lateral contacts.

**Figure 4.**
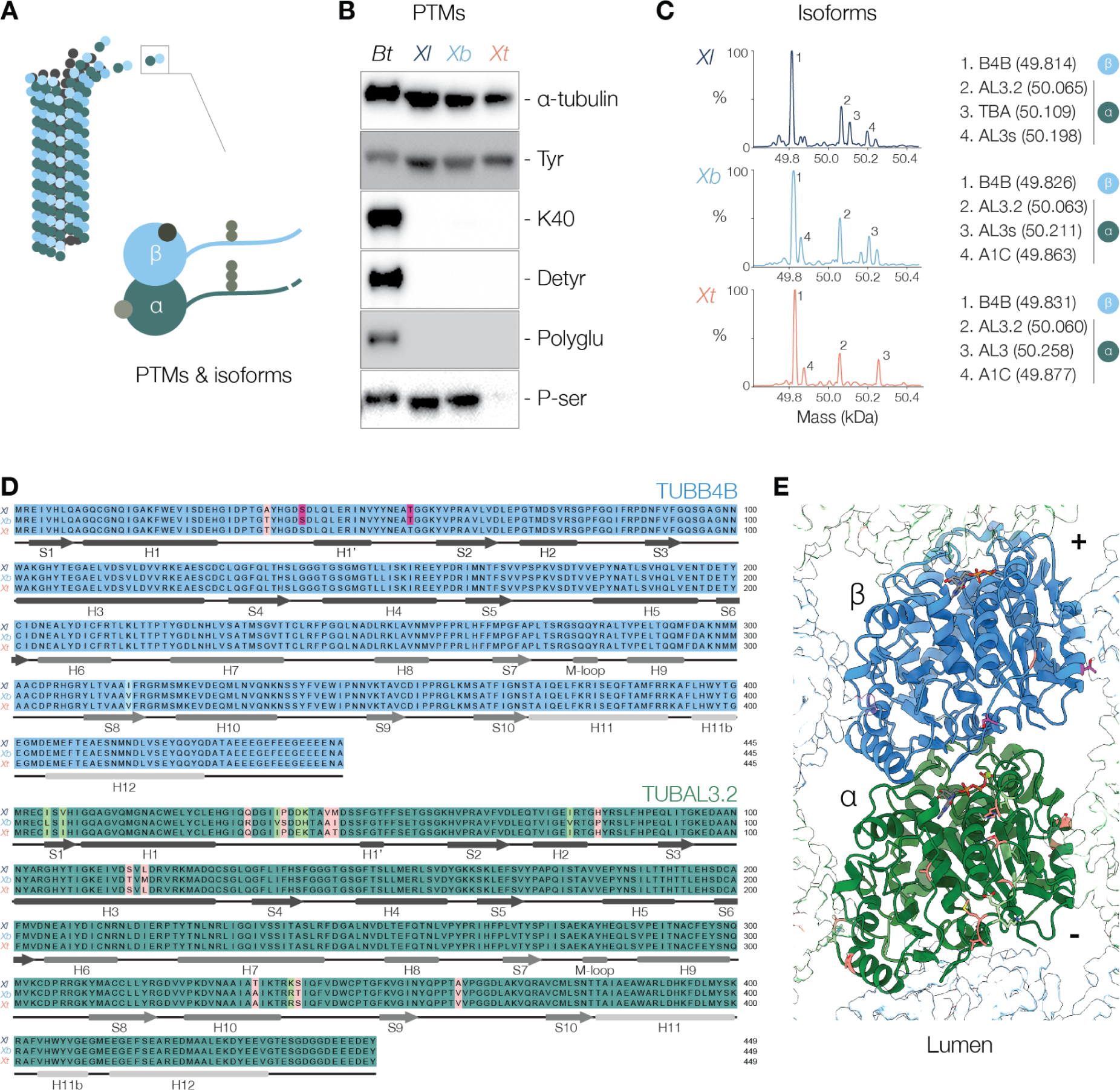
Sequence conservation between purified *Xenopus* tubulins (**A**) Microtubules can be composed of different tubulin isoforms and can carry posttranslational modifications (PTMs). (**B**) Western blot analysis of PTMs present in *B. taurus*, *X. laevis*, *X. borealis* and *X. tropicalis*. α-tubulin is a loading control, *B. taurus* is a positive control. Tyr binds with the last 8 residues of the C-tail of α-tubulin when tyrosinated. K40 recognizes the acetylation of α-tubulin on Lys residues. Detyr is a polyclonal antibody against detyrosinated α-tubulin. Polyglu reacts with polyglutamylated α- and β-tubulin. P-ser is a polyclonal antibody that recognizes proteins phosphorylated on serine residues. (**C**) Deconvoluted mass spectra of purified *Xenopus* tubulins. Measured average masses of the most abundant signals and corresponding tubulin isoforms. See also Figure S6A-C. (**D**) Sequence alignment for the most abundant α- and β-tubulin isoforms. Similar amino acid differences are highlighted in light blue or green with larger differences shown in salmon. Potential phosphorylation sites are labelled in magenta (see Figure S6D). Beneath the alignments arrows and tubes represent β-strands and α-helices within the model – these are coloured according to the domain architecture within tubulin monomers (dark grey – N-terminal domain; grey – intermediate domain; light grey – C-terminal domain). (**E**) Tubulin model for *Xl* with sequence differences to *Xb* and *Xt* mapped on according to colours from sequence alignment in (D). See also Figures S6.

### GDP-tubulin lattices are similar across the *Xenopus* species with subtle differences in the β-tubulin lateral interactions

The microtubule lattice is held together by longitudinal and lateral tubulin interactions. To investigate the relative contributions of these interactions, we aligned the 14-3 models by the N-terminal domain of a single β-tubulin monomer (denoted by an asterisk in Figure 5A, E, F) and coloured residues by the root-mean-square-deviation (RMSD) between species. When comparing the models of *X. laevis* and *X. tropicalis* – the most divergent species – we found the core of the tubulin monomers to be within 1Å of each other (Figure 5A, white) indicating similar longitudinal positioning along a protofilament. Consistently, the intra- and inter-dimer lattice spacing measurements were similar across all species and also maintained across different architectures. Further, the longitudinal lattice spacing (Figure 5B) and the full dimer rise measurements (∼ 81.1Å) are consistent with previous models for GDP-bound tubulin lattices,^41,43,44^ and with our cryo-EM density indicating GDP is bound to β-tubulin (Figure S5).

**Figure 5.**
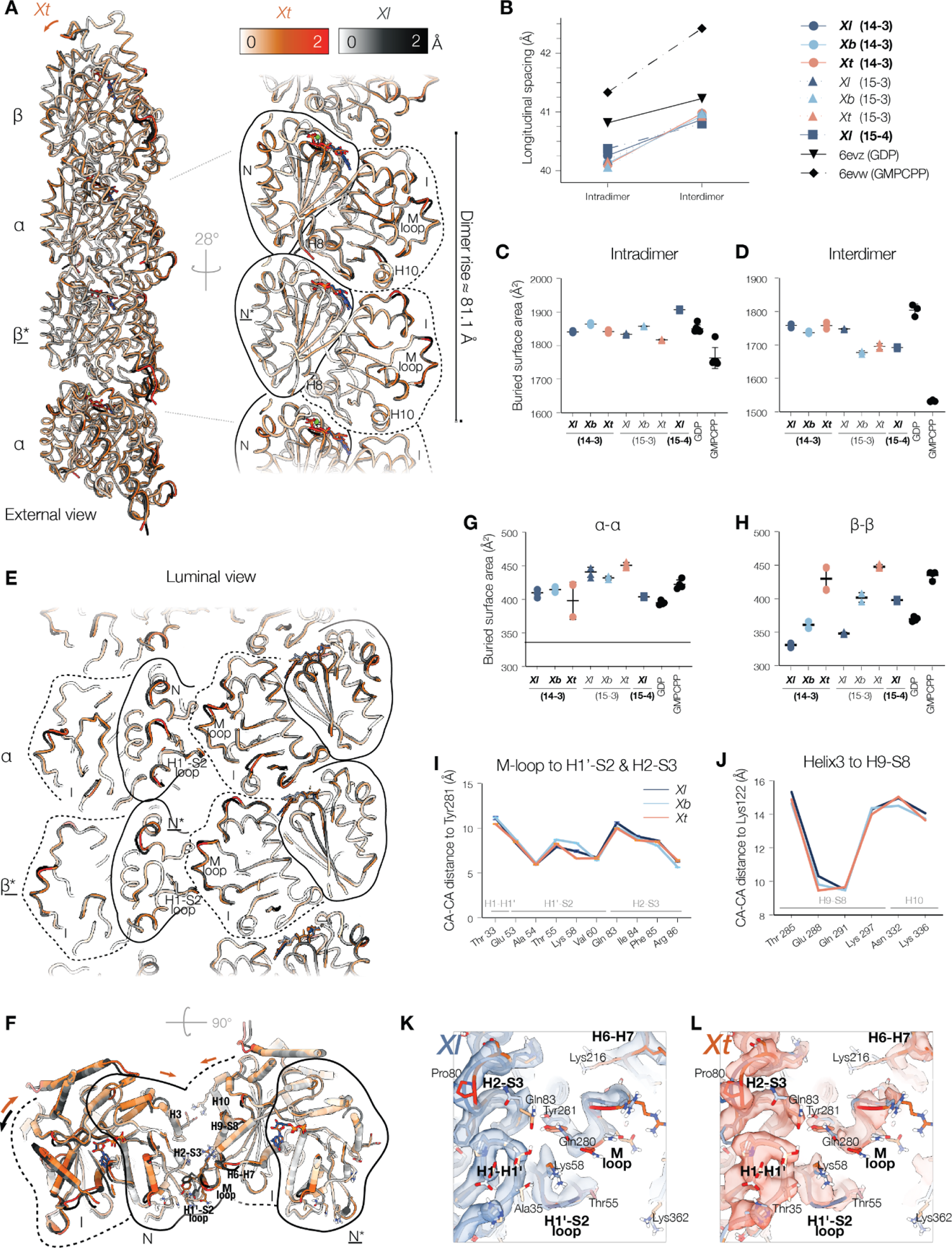
Tubulin lattice model analysis and comparison between isolated *Xenopus* tubulins with porcine tubulin in two different nucleotide states (**A**) Two tubulin dimers within a single protofilament from the models of 14 protofilament 3-start (14-3) microtubules from *X. laevis* and *X. tropicalis*. Models are coloured along a gradient according to the root-mean-squared deviation (RMSD) following alignment by the N-domain of a single β-tubulin (<u>N*</u>). No deviance is white and ≥2 Å deviance is coloured black for *Xl* and orange for *Xt*. (**B**) Intra- and interdimer α-β distance measurements reflective of longitudinal spacing within the tubulin lattice. Data is shown for 14 and 15 protofilament microtubules for each *Xenopus* species and compared with previously published porcine tubulin lattice reconstructions (black) containing either GDP (pdb – 6evz) or GMPCPP (pdb – 6evw).^44^ (**C-D**) Total α-β intradimer (C) and interdimer (D) protein-protein interface areas calculated for all *Xenopus* models compared with porcine tubulin models in two different nucleotide states as in (B). Determined by PISA analysis (M&M).^45^ (**E**) A cut through two neighbouring protofilaments within *X. laevis* and *X. tropicalis* models of 14-3 microtubules depicting the lateral interactions across two protofilaments aligned and depicted as in (A). (**F**) Cross section of β-tubulin from two neighbouring protofilaments within *X. laevis* and *X. tropicalis* models of 14-3 microtubules – aligned and depicted as in (A). (**G-H**) Total α-α tubulin (C) and β-β interdimer (D) lateral protein-protein interface areas calculated for all *Xenopus* models compared with porcine tubulin models in two different nucleotide states as in (B, C-D). Determined by PISA analysis (M&M).^45^ (**I-J**) Distance measurements between the Cα atoms of Tyr281 in the M-loop (K) and Lys122 in Helix3 with the Cα atoms of nearby residues within the lateral interfaces of neighbouring tubulins. (**K-L**) Aligned models and corresponding EM density for 14-3 microtubules at the lateral β-β tubulin interface for *X. laevis* (K) and *X. tropicalis* (L).

We next conducted Proteins, Interfaces, Structures and Assemblies (PISA) analysis of our models to quantify the predicted interactions between monomers.^45^ Longitudinally, we found that the interacting surface areas at the intra- and inter-dimer interfaces were (i) similar across all three *Xenopus* species, (ii) conserved between architectures, and (iii) similar to the porcine GDP-tubulin lattice (Figure 5 C, D, 6evz).^44^ Overall, our analysis found no major differences in the longitudinal interactions between *Xenopus* species.

Next, we analysed the lateral homotypic tubulin interactions. The RMSD between *X. laevis* and *X. tropicalis* indicated higher deviation at the lateral interface, particularly in the position of the M-loop, which differed slightly between species (Figure 5A, F, E). PISA analysis of lateral β-β tubulin interactions revealed the largest interface area for *X. tropicalis* and the smallest area for *X. laevis* across all helical architectures (Figure 5H). This was surprising given that neither of the two amino acid substitutions across β-tubulin (Figure 4D, E) are located at the lateral interface. To explore the structural origins of these differences, we measured distances between the α-carbons of key amino acids (Lys122 and Tyr281) and residues in the neighbouring β-tubulin at the lateral interface (as identified by PISA) (Figure 5 I, J). Overall, these indicated that backbone positioning of lateral contacts between species was very similar, and that the changes reflected in the PISA analyses were at the level of individual side chains. Although the resolution of our reconstructions did not provide precise information about the position of every residue, a qualitative inspection of side chain density indicated that there were some differences between species: in *X. tropicalis*, Gln83 was oriented toward the M-loop of the neighbouring tubulin, while in *X. laevis*, Gln83 was directed away from the lateral interface with a likely concomitant reduction in interaction strength (Figure 5 K, L).

Weakening of lateral contacts has been linked to the accumulation of lattice strain following GTP hydrolysis,^44^ but has also been proposed as a mechanism for increasing microtubule stability at sites of damage.^46–48^ Additionally, simulation data have suggested a link between loop ordering within the free tubulin dimer and the activation energy required for lattice incorporation.^49^ Thus, while the mechanistic link between structuring lateral loops, lateral interaction strength, and strain accumulation is unclear, it is reasonable to assume that a cumulative effect of changes in lateral contacts can contribute to differences in microtubule dynamics. Indeed, the observed 100 Å^2^ difference in β-β interaction surface area between *X. laevis* and *X. tropicalis* are on a similar scale as between GMPCPP and GDP models (6evw – 435.5 Å^2^; 6evz – 369.9 Å^2^) (Figure 5H). This indicated that although small, the scale of structural changes, particularly when cumulative, was sufficient to impact overall microtubule dynamics and stability. Our structural data suggested that *X. laevis* microtubules have the weakest lateral interactions (Figure 5H), while our dynamic data suggest that they have the most stable lattice (Figure 2), consistent with the idea that weakened lateral contacts can contribute to overall stability. As an indication of strain tolerance, we explored levels of lattice distortions by comparing microtubule models fit to symmetrised and non-symmetrised microtubule reconstructions.^50^ Comparison of 14-3 microtubules across species indicated that within our cryo-EM samples, both *X. laevis* and *X. borealis* microtubules contained a higher degree of lattice distortion whilst *X*. *tropicalis* microtubules are comparatively uniform (Movie S1). Thus, weaker lateral interactions in *X. laevis* could increase the tolerance for distortion or bending at local lattice contacts and contribute to their observed microtubule stability. Conversely, stronger lateral interactions in *X. tropicalis* could reduce tolerance for distortion contributing toward instability and higher catastrophe rates (Figure 1E, 2H).

Overall, the high sequence conservation of tubulin in the three *Xenopus* species is reflected in the overall structural similarity of their lattices, albeit with small differences at their β-tubulin lateral interactions. We therefore set out to explore more specific differences in their dynamics at the growing microtubule plus-end.

### Tubulin’s apparent activation energy scales with ambient temperature

GTP hydrolysis is essential to microtubule dynamics as it determines the size of the protective GTP-cap (Figure 6A).^26,51–53^ The microtubule-binding protein EB1 recognizes the nucleotide state of growing microtubule ends and thus serves as a proxy of the GTP-cap size and hydrolysis rates.^50,54–57^ We equalised the growth rates of all three microtubule species, imaged purified GFP-tagged EB1, and used sub-nanometer tracking and comet averaging to measure the EB1 comet length in all three *Xenopus* species (Figure 6B). As reported previously,^53^ cap size increased with growth velocity (Figure 6C). Fitting the averaged EB1 profiles gave a comet length of λ = 185 ± 1 nm for *X. laevis*, λ = 224 ± 1 nm for *X. borealis* and λ = 201 ± 1 nm for *X. tropicalis* (Figure 6D). We concluded that there was no significant difference in the GTP hydrolysis rate, which was supported by almost no divergence around the nucleotide binding pockets (Figure 6E). We next determined the activation energy of the forward reaction of linear polymer growth via an Arrhenius plot (Figure 6F). Here, the activation energy (E_a_) can be derived from the slope of ln(k) versus 1/T with E_a_ (*Xl*) = 41 ± 2 kJ mol^-1^, E_a_ (*Xb*) = 47 ± 8 kJ mol^-1^, and E_a_ (*Xt)* = 60 ± 7 kJ mol^-1^. Interestingly, the cold-adapted *Xenopus* tubulin had the lowest activation energy and the warm-adapted tubulin the highest activation energy. In a simple model of linear microtubule polymer growth,^58^ this means that *X. laevis* tubulin has the highest free energy in solution and thus incorporates more easily into the growing microtubule plus-end. Similarly, fast microtubule growth in *C. elegans* has recently been explained by a more active tubulin dimer.^49^ Taken together, the inverse scaling of the tubulin’s free energy and the decrease in flexibility of lateral contacts with temperature provide an explanation for the higher microtubule growth rates, the low critical concentration, and the overall stability of microtubules in cold-adapted frog species (Figure 6G).

**Figure 6.**
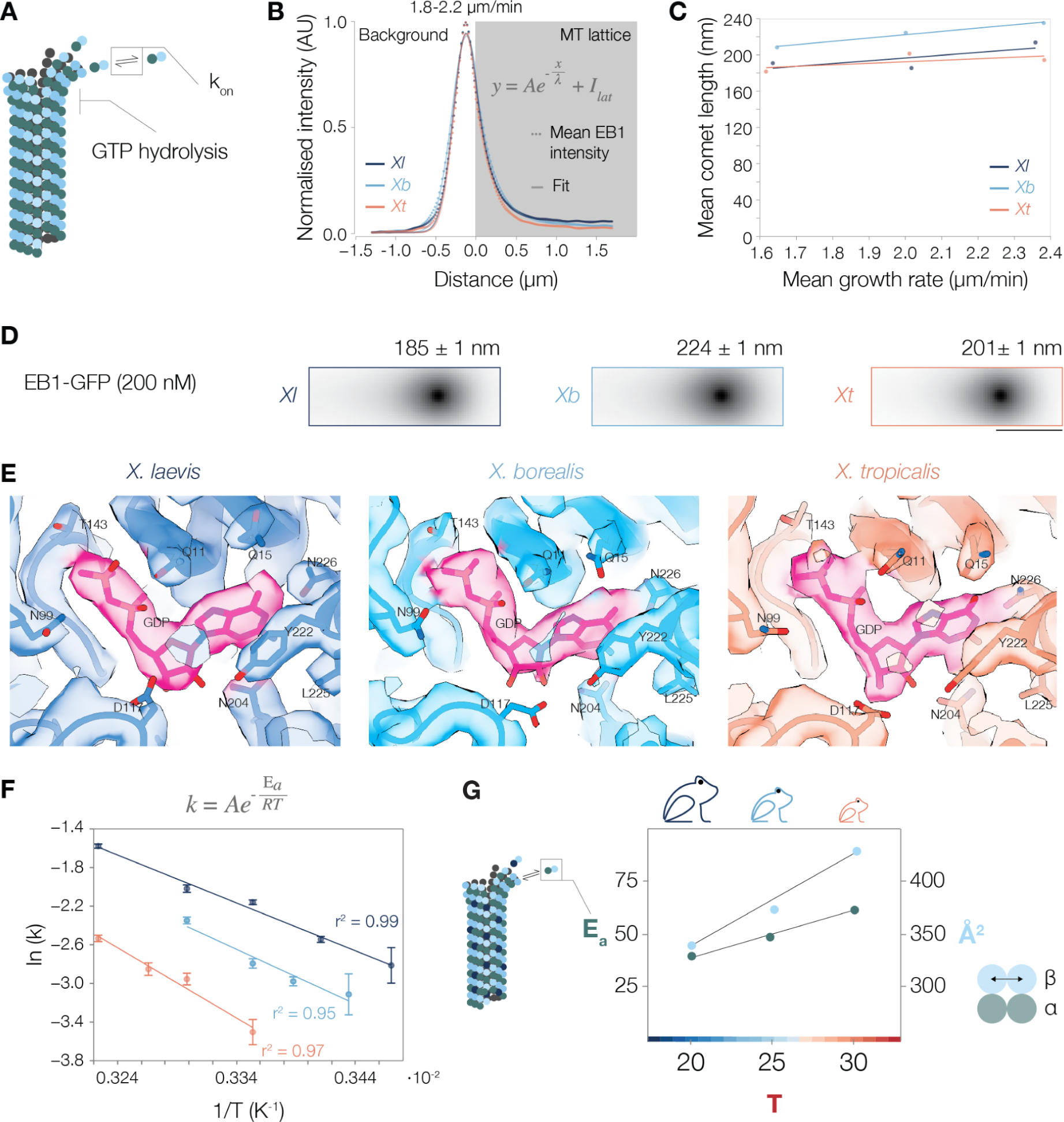
Tubulin’s apparent activation energy scales with ambient temperature (**A**) At the plus-end, microtubule growth rate reflects a balance of GTP-tubulin association and dissociation. Away from the GTP-cap, GTP is hydrolysed to GDP driving dynamic instability. (**B-D**) Analysis of EB1-GFP profiles from microtubules growing at 25°C, with 200 nM EB1-GFP and tubulin concentration 12 µM for *Xl*, 11-15 µM for *Xb*, 25-28 µM for *Xt* to adjust for similar growth velocities. (**B**) Plot of super average of EB1-GFP intensity profiles centred around peak intensity. Microtubule lattice corresponds to the curve with a grey background, which was fitted with an exponential decay function (starting at 0 which is defined one pixel after the peak intensity). The rest of the curve was fitted with a Gaussian function. Each point is the average mean intensity for a subpixel, error bars are SEM. (**C**) Average mean comet length (λ), obtained by binning EB1-GFP profiles with growth velocity (1.4-1.8, 1.8-2.2 and 2.2-2.6 µm/min) and fitting an exponential decay function as shown in (B). At 1.4-1.8 µm/min: v_g,*X*l_=1.64 ± 0.02 µm/min, λ*_Xl_*=190 ± 2 nm (197 profiles); v_g,*Xb*_=1.65 ± 0.02 µm/min, λ*_Xb_*=208 ± 1 nm (448 profiles); v_g,*Xt*_=1.62 ± 0.01 µm/min, λ*_Xt_*182 ± 1 nm (645 profiles). At 1.8-2.2 µm/min: v_g,*Xl*_=2.02 ± 0.01 µm/min, λ*_Xl_*=185 ± 1 nm (1223 profiles); v_g,*Xb*_=2.00 ± 0.01 µm/min, λ*_Xb_*=224 ± 1 nm (1091 profiles); v_g,*Xt*_=2.01 ± 0.01 µm/min, λ*_Xt_*=201 ± 1nm (581 profiles). At 2.2-2.6 µm/min: v_g,*Xl*_=2.36 ± 0.01 µm/min, λ*_Xl_*=213 ± 2 nm (481 profiles); v_g,*Xb*_=2.38 ± 0.02 µm/min, λ*_Xb_*=235 ± 2 nm (643 profiles); v_g,*Xt*_=2.38 ± 0.02 µm/min, λ*_Xt_*=194 ± 2 nm (238 profiles). v_g_: mean ± SEM, λ: mean ± SE. Linear regression shows ordinary least squares. (**D**) Average EB1-GFP signal from the sub-pixel (6-time subsampling) alignment of comets peak intensities (1223 profiles for *Xl*, 1091 for *Xb*, and 581 for *Xt*) at a growth velocity of 1.8 – 2.2 µm/min. Scale bar: 500 nm. Avg. length +/- SE is shown. (**E**) EM density and models at the site of GTP hydrolysis within β-tubulin for 14 protofilament 3-start microtubules for *Xl*, *Xb*, and *Xt*. (**F**) Arrhenius plot of the tubulin polymerization reaction for *Xl*, *Xb*, and *Xt*. k values were obtained from the plots of Figure 2E-G. The log-function ln(k)=ln(A)-E_a_/RT was fitted to the average k using ordinary least square regression. Error bars show standard error. (**G**) Conceptual model for the mechanistic basis of the observed differences in microtubule dynamics between three *Xenopus* species.

## Discussion

Tubulins are among the most conserved proteins. This high degree of conservation is particularly evident between different *Xenopus* species, where tubulin sequences are remarkably similar (Figure 4D). Therefore, it was surprising to observe significant differences in microtubule dynamics between these species. This, however, makes purified *Xenopus* tubulins an excellent tool for investigating how subtle differences in primary sequence can biochemically and structurally explain micron-scale variations in microtubule dynamics. In this study, we found cold-adapted tubulin to exhibit a lower activation energy and thus incorporate more readily into the growing microtubule plus-end. This characteristic can explain the increased growth velocities and lower critical concentrations observed at lower temperatures. Structurally, we found the microtubule lattice of cold-adapted tubulin to be more flexible in its lateral contacts, reducing the strain that typically accumulates in the structure. This flexibility likely contributes to the overall stability of the microtubules under cold conditions. While the observed structural differences are subtle, it is important to realise that even slight variations accumulate in a microtubule, which is several micrometres long and composed of thousands of tubulin subunits.

Dynamic instability is an intrinsic property of microtubules, which means that changes in tubulin’s primary sequence can directly modulate dynamics. Indeed, *in vitro C. elegans* microtubules combine fast growth rates with frequent catastrophes,^49^ while human microtubules combine fast growth with longer lifetimes.^59,60^ This implies that catastrophe frequency is not simply a function of microtubule growth. In this study, we purified tubulins from three *Xenopus* species that showcase such combinatorial variety in microtubule growth and catastrophe frequency. Adjusting the temperature close to the respective ambient temperature equalised polymerization velocity across the three microtubule species (Figure 1C). The catastrophe frequencies, however, remained significantly different (Figure 1E). Thus, while the general assumption that faster growing microtubules are stabilised by their larger GTP-caps appears to be a conserved mechanism,^26,51–53^ catastrophe frequency might be determined on a species level by modulating the strength of lateral contacts between neighbouring protofilaments as suggested by our structural data (Figure 5).

In a cellular context, microtubule dynamics are controlled by a variety of MAPs and motors. The γ-tubulin ring complex (γ-TuRC), for example, is the principal microtubule nucleation template that gives rise to canonical 13-protofilament microtubules.^61–63^ For structural analysis, the low critical concentration of *Xenopus* tubulins (Figure 1D) allowed us to grow the microtubules in the absence of microtubule-stabilising agents or seeds. Therefore, the observed differences in architectural distribution (Figure 3D) must ultimately have arisen as a consequence of tubulin interactions specific for each *Xenopus* species. Indeed, recent studies revealed that during nucleation, the dynamic assembly of tubulin into microtubules occurs in multiple sequential steps with different lateral tubulin interactions formed in different intermediates.^64^ Thus, the different lateral interactions we observe within the tubulin lattices could impact microtubule nucleation dynamics *in vivo*.^65^

Similarly, plus-end dynamics are modulated in a cellular context, where microtubule growth velocity and catastrophe frequency mainly determine microtubule length distribution and thus final polymer mass. Indeed, we understand how changes in proteins that regulate microtubule dynamics can control spindle size,^66–69^ which in case of our three *Xenopus* species scales with cell and body size.^70,71^ Our data explain how tubulin, as the spindle’s main building block, can contribute to the control of microtubule mass and therefore steady-state spindle length. Therefore, microtubule dynamics are unlikely to be governed by MAPs and motors alone – to gain a complete understanding, we need to appreciate the contribution of tubulin’s intrinsic properties. Thus, while the control of microtubule dynamics by MAPs and motors might exert a selective pressure on tubulin sequence diversification^72^, subtle and allosteric changes are essential to divergent mechanisms that convey specialised function.

## SUPPLEMENTARY INFORMATION

**Supplementary Figure 1:**
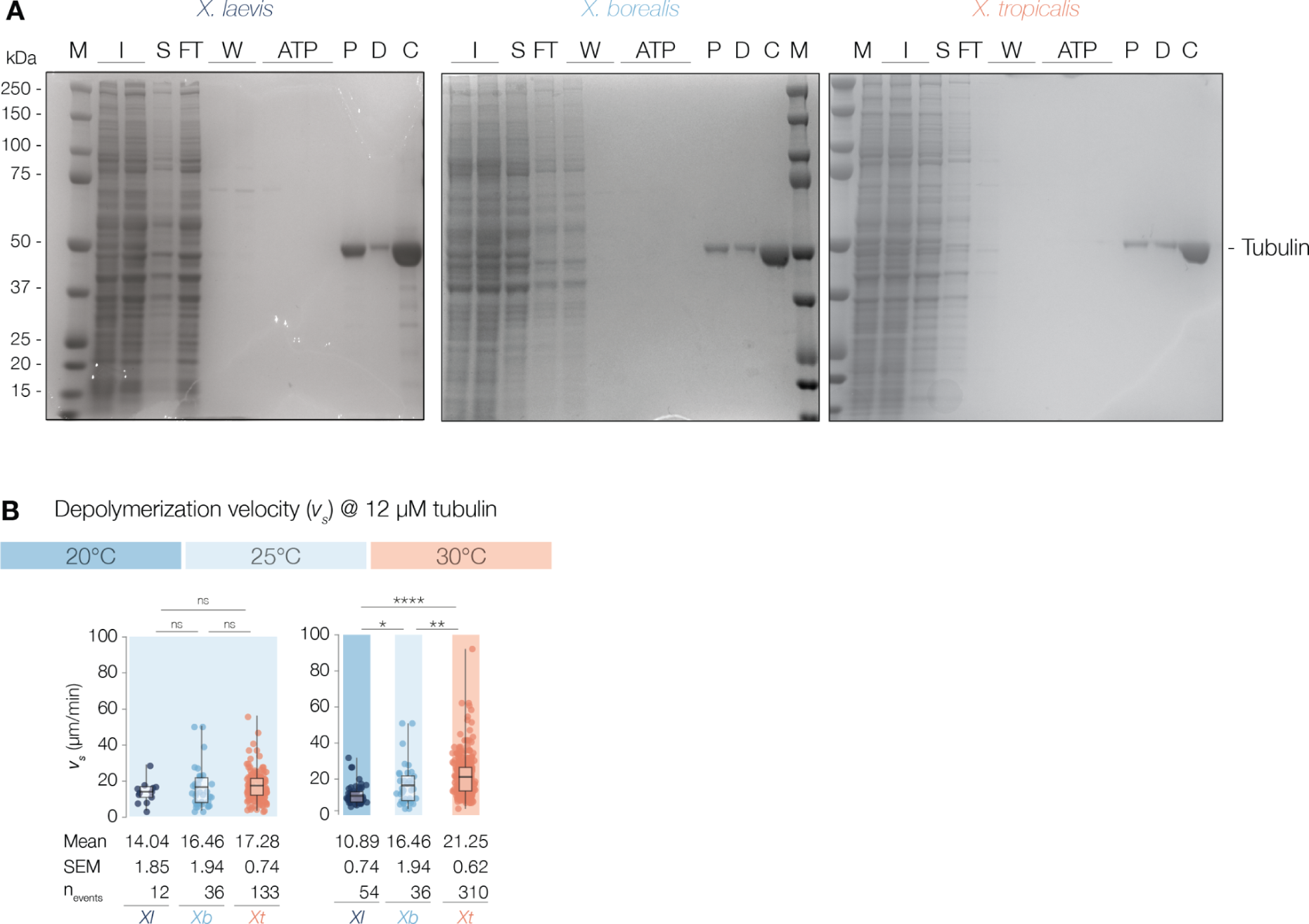
Tubulin purification from extracts prepared from unfertilized *X. laevis*, *X. borealis*, and *X. tropicalis* eggs, related to Figure 1. (**A**) Coomassie-stained SDS-PAGE of tubulin purifications. M: Marker, I: Input, S: Supernatant, FT: Flowthrough, W: Wash, ATP: ATP wash, P: Peak fraction, D: Desalt, C: Concentrated. (**B**) At 25°C, *Xl* microtubules depolymerize at 14.04 ± 1.85 µm/min, *Xb* at 16.46 ± 1.94 µm/min, *Xt* at 17.28 ± 0.74 µm/min, with p-value(*Xl,Xb*)=0.88, g(*Xl,Xb*)=0.22; p-value(*Xb,Xt*)=0.18, g(*Xb,Xt*)=0.09; p-value(*Xl,Xt*)=0.21, g(*Xl,Xt*)=0.38. Right graph (20°C *Xl*, 25°C *Xb*, 30°C *Xt*): p-value(*Xl,Xb*)=0.03, g(*Xl,Xb*)=0.65; p-value(*Xb,Xt*)<0.01, g(*Xb,Xt*)=0.43; p(*Xl,Xt*)<0.0001, g(*Xl,Xt*)=1.00. All values were obtained from measurements of microtubules pooled over at least three independent experiments, all p-values were calculated with the Mann-Whitney test, g corresponds to Hedge’s g. Data points correspond to individual measurements. All error values are SEM. Plot boxes range from 25th to 75th percentile, whiskers span the range, horizontal line shows median value.

**Supplementary Figure 2:**
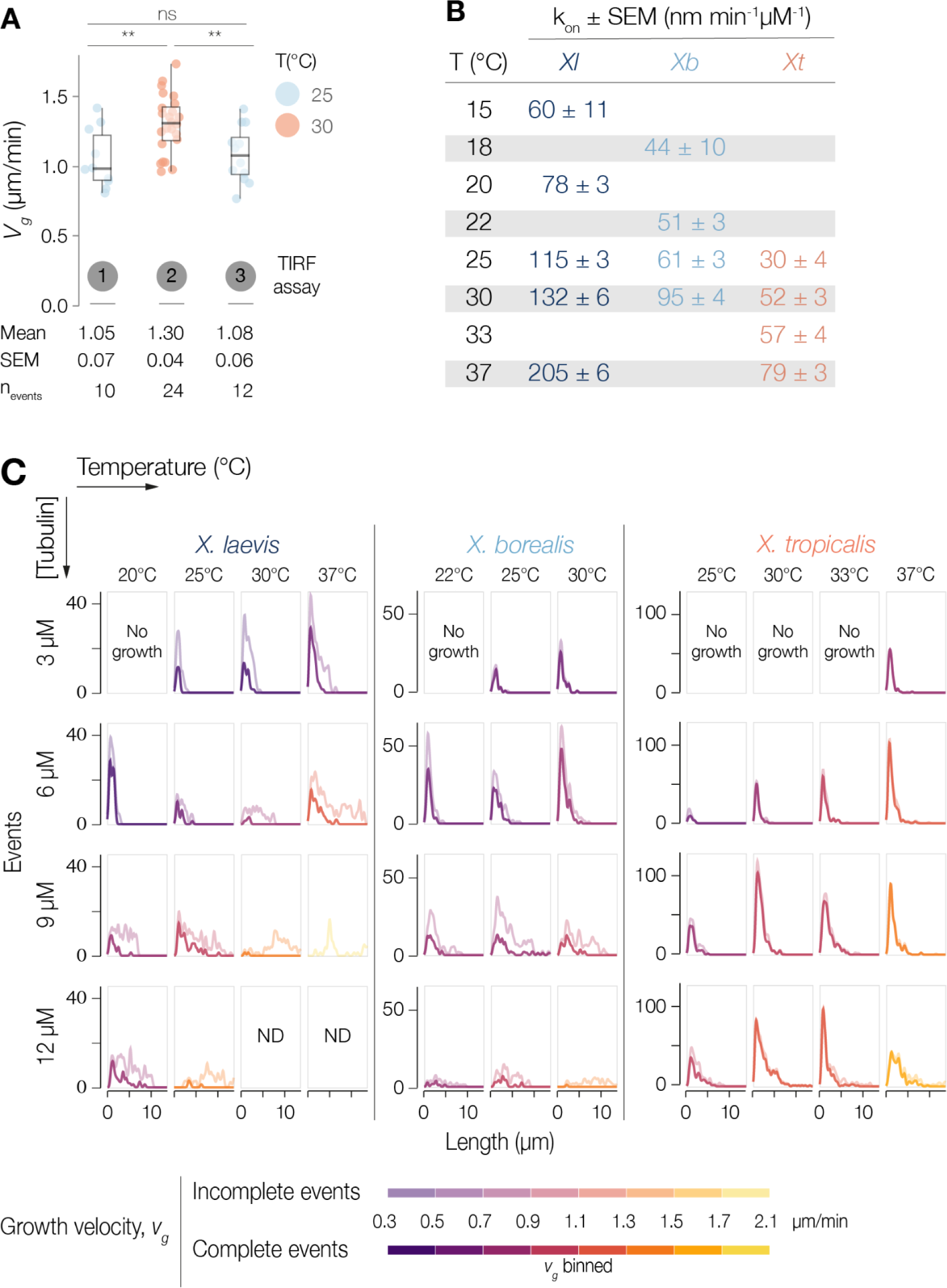
*X. laevis* and *borealis* microtubules switch to unbounded growth at higher growth rates, related to Figure 2. (**A**) Example of growth velocity measurements using three consecutive 10-min acquisition intervals at different temperatures: (1): 25°C, (2): 30°C, (3): 25°C. Mean difference between intervals (1) and (3) is not significant (p-value(1-3)=0.71, hedge’s g(1-3)=0.15). p-value(1-2)<0.01, hedge’s g(1-2)=1.18. p-value(2-3)<0.01, hedge’s g(2-1)=1.04. p-values tested with Mann-Withney. (**B**) Slopes of the regressions from Figure 2E-G, in nm.min^-1^.µM^-1^. (**C**) The probability distribution of microtubule length is stacked for complete (opaque) and incomplete (transparent) events and pooled per temperature, tubulin concentration, and species. Microtubule lengths are 2 μm-binned (0-18 μm). For each condition, the average growth velocity is calculated and conditions are colour coded by growth velocity intervals. ND = not determined.

**Supplementary Figure 3:**
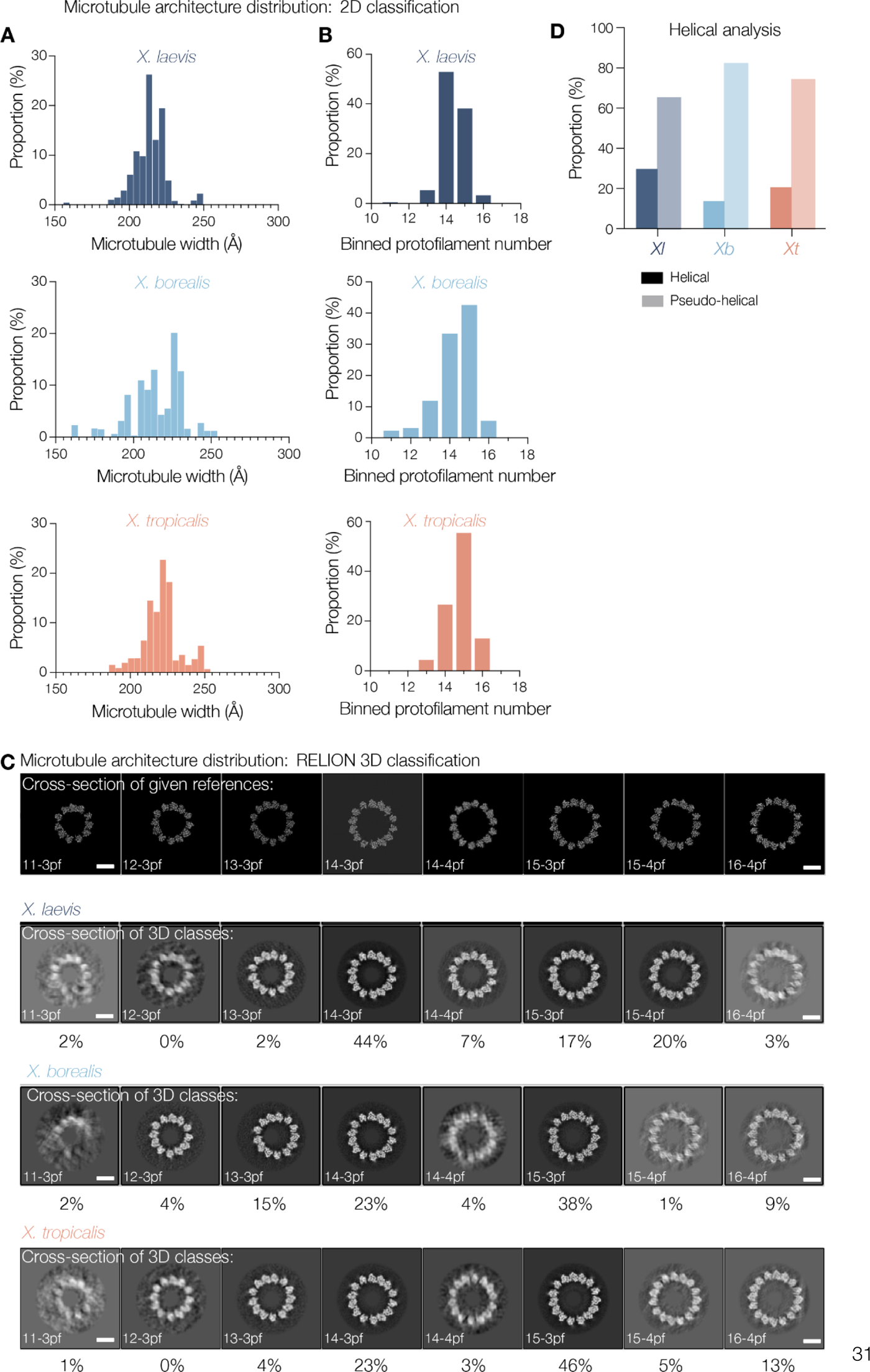
Architectural distribution as determined by RELION 2D and 3D classification, related to Figure 3. (**A**) Histograms showing the width distribution of the particles within the top 120 classes following RELION 2D classification. Width is binned by pixel size (M&M). (**B**) Data binned by protofilament number according to width measurements of previously published reconstructions of 11-16 protofilament microtubules.^74^ (**C**) Cross-sections showing the references used to seed the 8 class RELION 3D classification (top) and cross-sections representative of the data grouped into each class (bottom). Percentages denote the proportions of total particles assigned into each class. References were created using previously determined helical parameters.^42,74^ (**D**) Bar graph representing microtubule distributions from (C) grouped by helical start number reflective of a fully helical microtubule lattice (with 4-start helix) or pseudo-helical microtubule lattices containing a seam (3-start helix) – as depicted in Figure 3A.

**Supplementary Figure 4:**
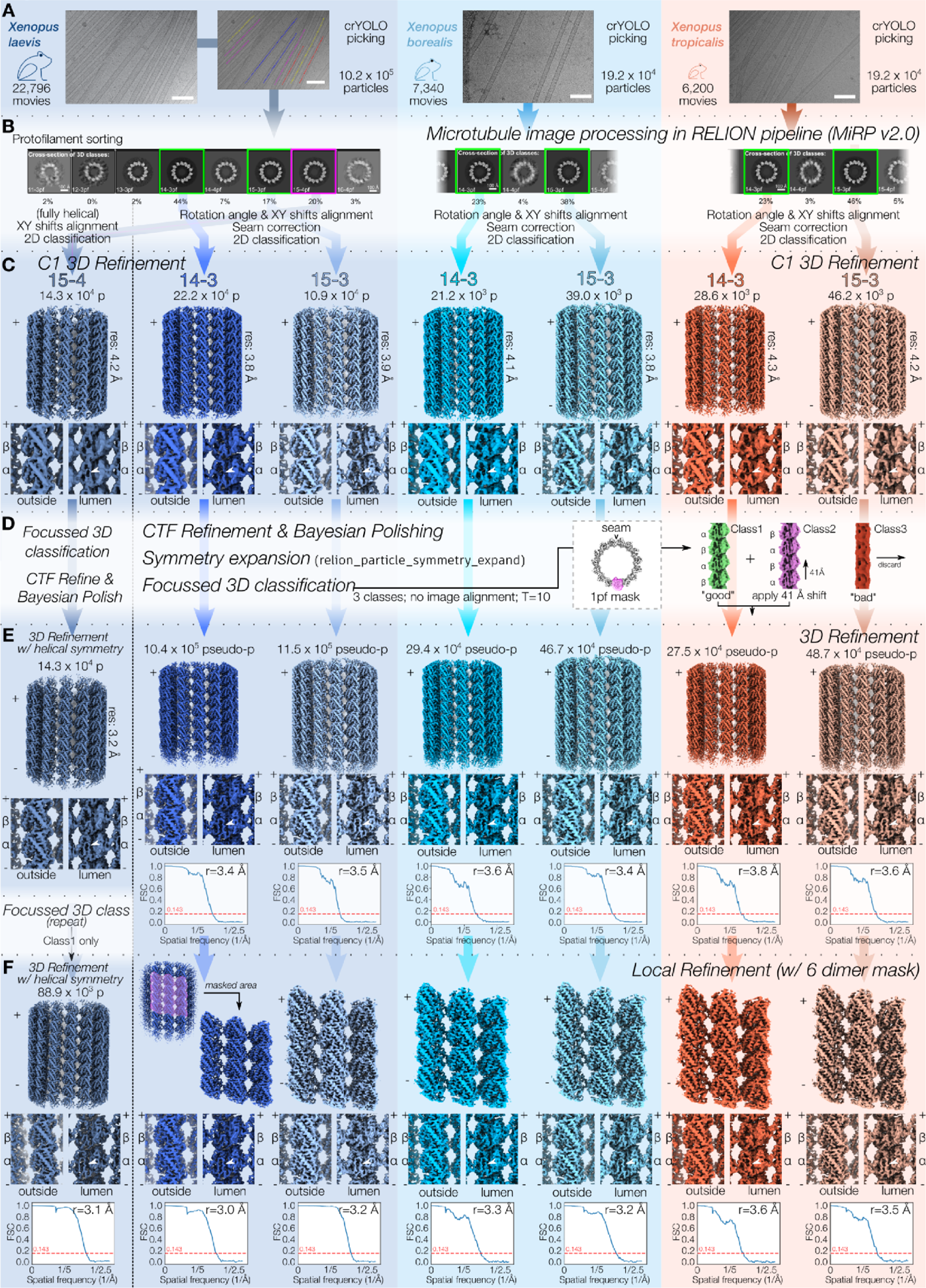
Microtubule cryo-EM image processing pipeline, related to Figure 3 and 5. (**A**) Exemplar micrographs for the cryo-EM microtubule dataset from each species, together with information about dataset size. Additionally, an example of crYOLO particle picking (coloured dots) of the *X. laevis* microtubule micrograph is shown. (**B**) 3D classification of picked particles enables protofilament architecture distribution in each dataset. For all three species, particles with 14-3 and 15-3 architectures (green boxes) were treated similarly. Particles classified as derived from fully helical (15-4 microtubules from *X. laevis*, pink) were treated differently because seam alignment/correction steps were not required, shown on the far left. (**C**) Initial rounds of processing to calculate C1 3-dimensional reconstructions for each architecture (top). The appearance of the S9-S10 loop density in α-tubulin opposite the seam (bottom panel, white arrowhead) is indicative of particle alignment accuracy, and in particular of accurate seam alignment. At this stage in the reconstruction process, the S9-S10 loop density is visible but discontinuous and requires further optimisation. (**D**) Individual protofilaments from 3-start microtubules were further processed through iterations of RELION Bayesian Polishing and CTF refinement, symmetry expansion, and focussed 3D classification of individual protofilaments. An indicative output of the protofilament 3D classification step is shown on the right, with pseudo-particles separated into three (Class 1 – well aligned; Class2 – one tubulin monomer out of register; Class3 – ‘junk’). For helical 15-4 microtubules: the same focussed 3D classification was used to correct for the register of the microtubules (with CTF refinement and Bayesian polishing), but symmetry expansion was not applied. Instead symmetry was exploited through imposition of helical symmetry within RELION 3D refinement. (**E**) Symmetry expanded 3-dimensional reconstructions showed substantial improvements in density quality (top), discrimination between α- and β-tubulin (middle, white arrowhead indicates S9-S10 loop density in α-tubulin) and overall resolution as indicated by the Fourier shell correlation (FSC) plots of two unfiltered half maps at a cutoff of 0.143 (bottom). For helical 15-4 microtubules, the application of symmetry constraints substantially improved the density quality. A final focussed 3D classification was repeated with only well-aligned particles assigned to class1 taken for the final reconstruction. (**F**) Final reconstruction from refinement of the helical lattice or 6 tubulin dimers opposite the seam shows further improvements in density (top), discrimination between α- and β-tubulin (middle, white arrowhead indicates S9-S10 loop density in α-tubulin) and overall resolution (bottom), all of which fall between 3.0 and 3.6 Å.

**Supplementary Figure 5:**
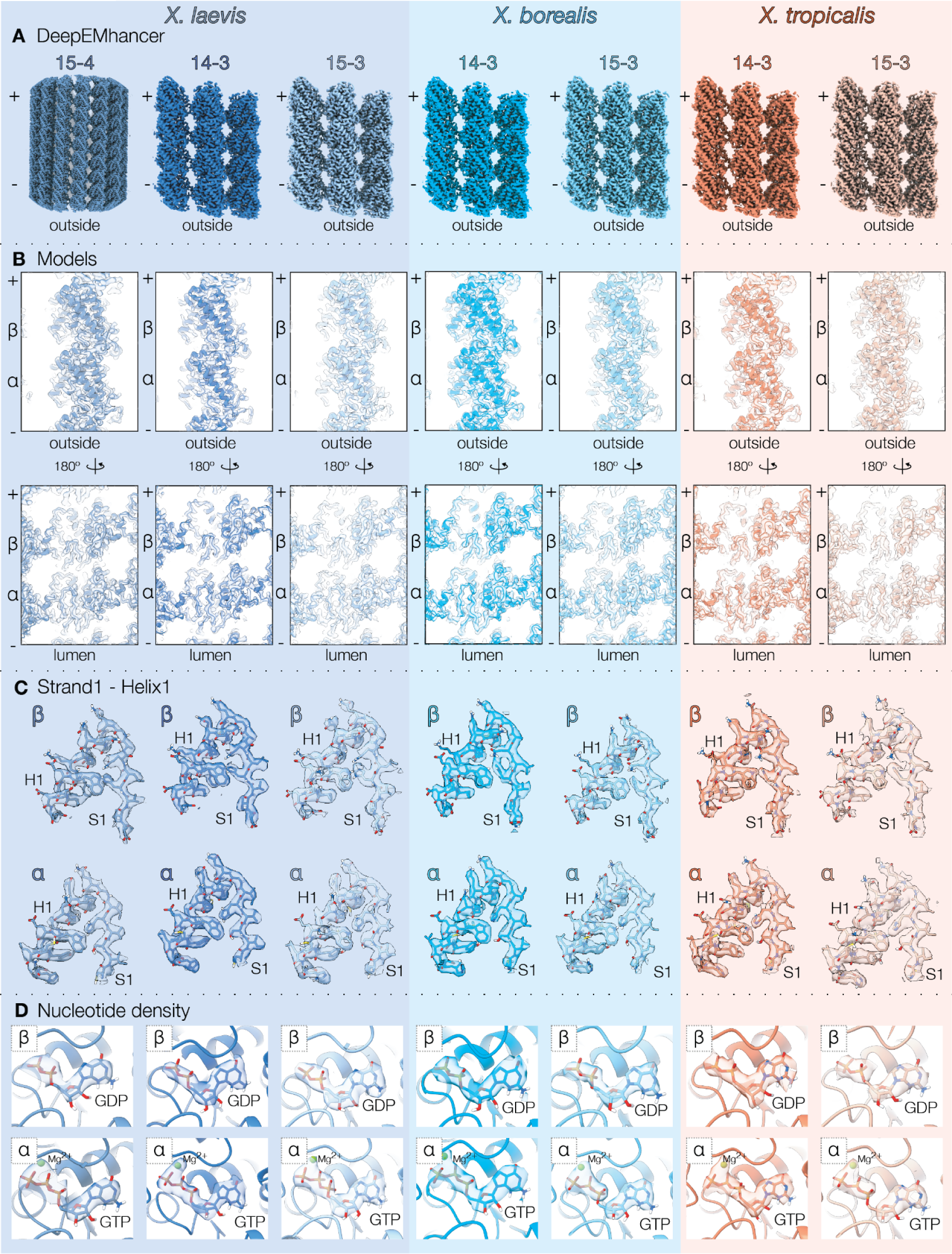
Cryo-EM post-processing and modelling, related to Figure 3. (**A**) Cryo-EM reconstructions for each architecture and species following the automated post-processing of masked half-maps by DeepEMhancer.^75^ This improved the density interpretability for model building (M&M). (**B**) A cut-through of the models within the DeepEMhancer processed density as viewed from outside the microtubule (top) or from the lumen (bottom). (**C**) Strand1-Helix1 (S1-H1) in β- and α-tubulin from our models shown within post-processed (DeepEMhancer) maps masked to this area to illustrate the quality of each reconstruction and fit of our models. (**D**) Density corresponding to the bound nucleotide within β-tubulin (top) and α-tubulin (bottom) reflective of the GDP and GTP-Mg^2+^ bound at each site.

**Supplementary Figure 6:**
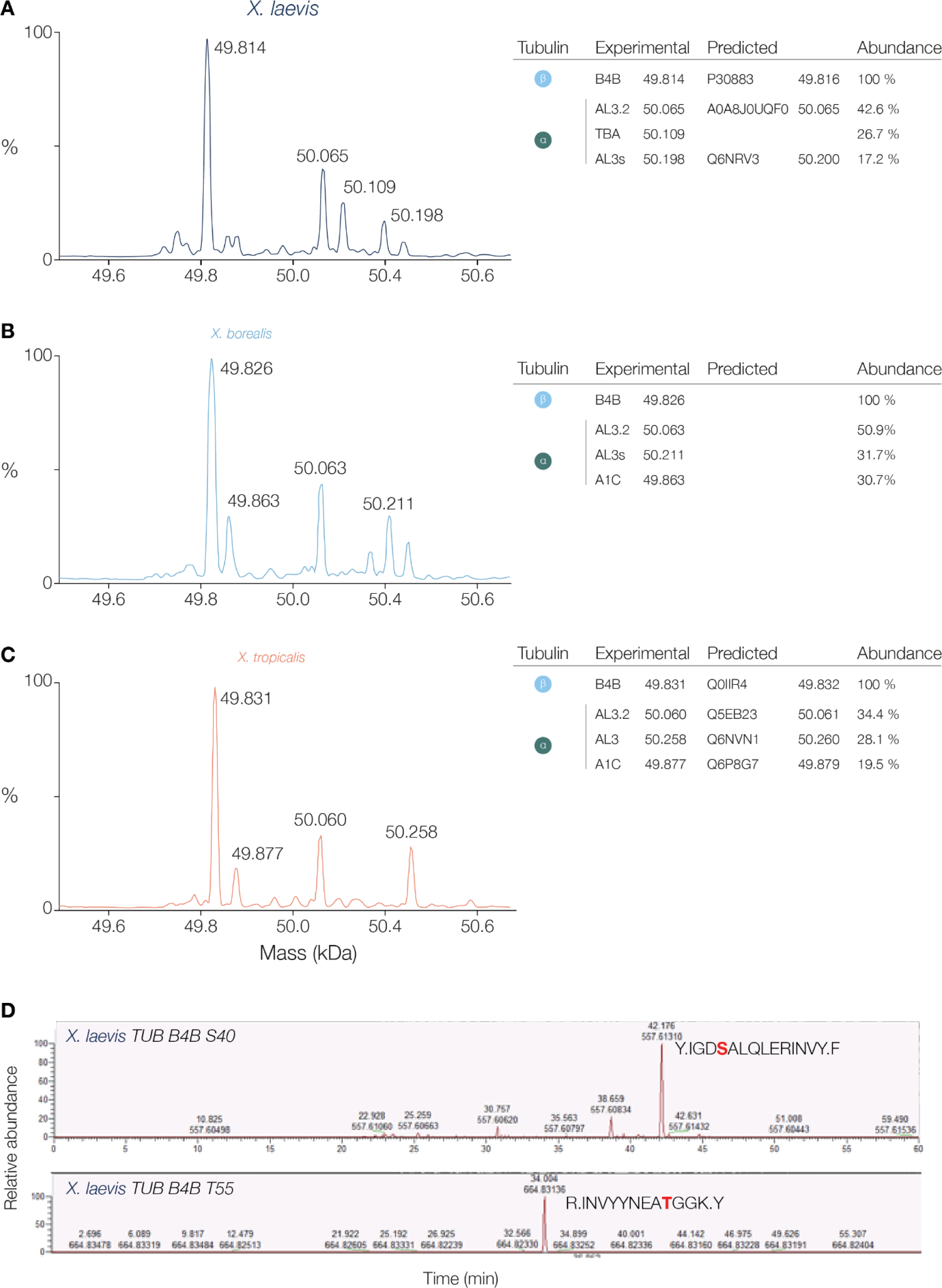
*Xenopus* tubulin isoforms and phosphosites as identified by mass spectrometric analyses, related to Figure 4. (**A-C**) Deconvoluted mass spectra of purified *Xenopus* tubulins. Measured average masses of the most abundant signals and corresponding tubulin isoforms are labelled on the right. Measured masses are in agreement with the theoretical masses of *Xl* B4B (MW=49.814 Da, Uniprot: P30883), *Xl* AL3.2 (MW=50.065 Da, Uniprot: A0A8J0UQF0), *Xl* TBD (to be determined), *Xl* AL3s (MW=50.198 Da, Uniprot: Q6NRV3), *Xt* B4B (MW=49.831 Da, Uniprot: Q0IIR4), *Xt* AL3.2 (MW=50.060 Da, Uniprot: Q5EB23), *Xt* AL3 (MW= 50.258 Da, Uniprot: Q6NVN1), and *Xt* A1C (MW=49.877 Da, Uniprot: Q6P8G7). For *X. borealis*, although the genome is fully sequenced, there are no annotations that include genes, transcripts or tubulin protein sequences. Peak assignment as described in M&M. (**E**) *X. laevis* tubulin was purified and phosphorylation enrichment was performed using an anti-pSer antibody. Samples were separated by SDS-PAGE and analysed by LC-MS/MS.

SUPPLEMENTARY MOVIE S1: Complete microtubule lattice distortion analysis

RMSD representations of complete 14 protofilament 3-start microtubule lattice models constructed from symmetrised and non-symmetrised cryo-EM density (frames alternate between the two models). A higher RMSD value (red) indicates a higher level deviance between the aligned microtubule particles and the perfectly cylindrical reconstructions generated through application of symmetry restraints. Higher deviance (red) reflects higher levels of lattice distortion within the data from each species.

**SUPPLEMENTARY TABLE S1:**
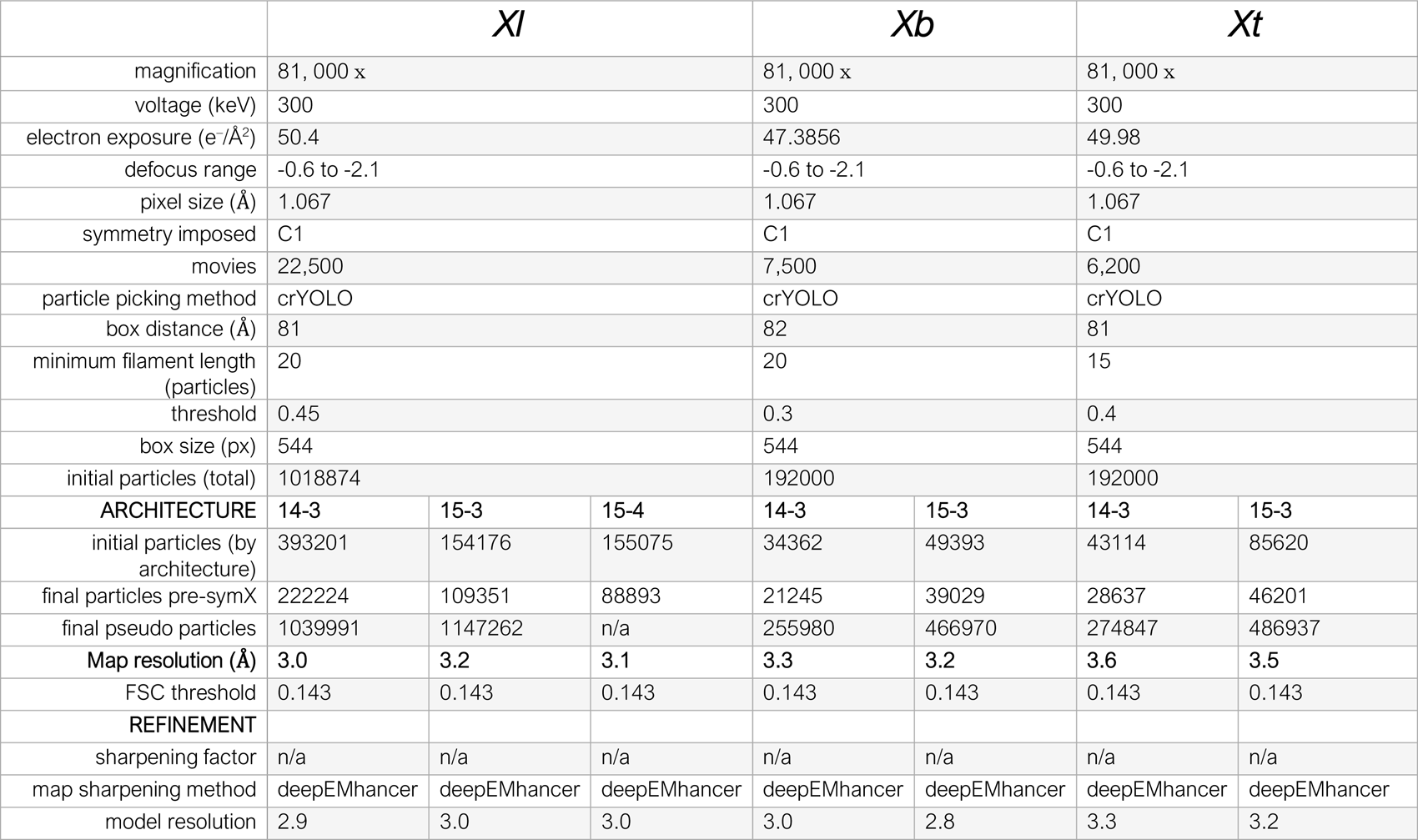
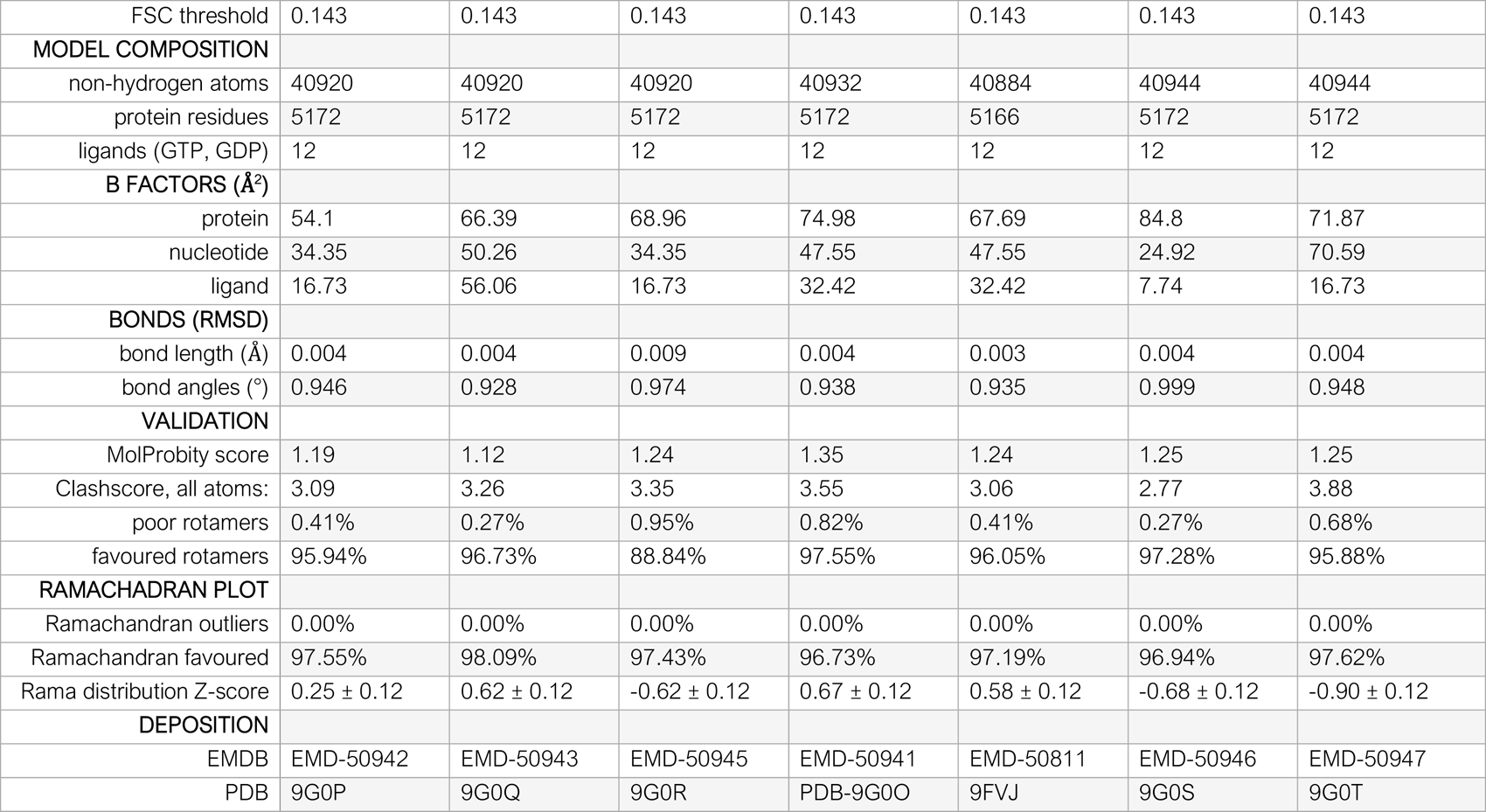
Cryo-EM data acquisition table.

## MATERIALS & METHODS

### Experimental model

*Xenopus* frogs (adult females obtained from NASCO, Fort Atkinson) used for this study were maintained at the animal husbandry of the Humboldt University in Berlin, in a recirculating tank system with monitored temperature (18-20°C for *Xl*, 19-23°C for *Xb*, 24-27°C for *Xt*) and water quality. They were fed with food pellets (V7106-0202) from ssniff Spezialdiäten GmbH. All experimental protocols involving frogs were performed in accordance with national regulatory standards and ethical rules, reviewed and approved by the LaGeSo under Reg.-Nr. 0113/20.

### Antibodies

Monoclonal anti α-tubulin (Sigma T5168) was used for PTMs identification and to assess the efficiency of tubulin purification. To map the PTMs present in different tubulins, following antibodies were used: Tyr: rat monoclonal (YL1/2, Abcam ab6160), recognizes the last 8 residues at the C-tail of α-tubulin when tyrosinated; K40: mouse monoclonal (Sigma T7451, clone 6-11 B-1), recognizes acetylation of α-tubulin on Lys residue; Detyr: rat polyclonal (Abcam ab48389), recognizes the C-terminal domain of detyrosinated α-tubulin; Polyglu: mouse monoclonal (AdipoGen GT335), reacts with polyglutamylated α- and β-tubulin; P-ser: rabbit polyclonal (Abcam ab9332), recognizes proteins phosphorylated on serine residues. Horseradish Peroxidase (HRP) antibodies anti-rabbit IgG (Proteintech 0001-2), anti-mouse IgG (Proteintech 0001-1) and anti-rat IgG (Invitrogen 62-9520) from goat were used as secondary antibodies.

### Tubulin purification

*Xl*, *Xb*, and *Xt* eggs arrested in metaphase of meiosis II were used to prepare Cytostatic factor (CSF) extracts from which tubulin was purified as previously described.^20^ Briefly, eggs were collected, lysed and spun in CSF-XB containing Cytochalasin D. The CSF-extract was diluted 1:1 in BRB80 buffer (80 mM PIPES, 1 mM EGTA, 1 mM MgCl₂, pH 6.9), and centrifuged for 10 min at 80.000 rpm in an MLA-80 rotor (Beckman-Coulter) at 4°C. The supernatant was loaded onto a TOG column and tubulin was eluted using high salt. Peak fractions were pooled, and buffer was exchanged to BRB80 supplemented 10 mM Mg^2+^-GTP using PD10 desalting columns (GE Healthcare). Protein was concentrated using a 30 kDa cut-off concentration filter column (Amicon), aliquoted in 5-10 µL aliquots and flash frozen in liquid nitrogen. Concentration was determined by measuring A_280_ values with a Nanodrop spectrophotometer (Thermo) after thawing and ultracentrifugation for 10 min at 2°C using a TLA-100 rotor (80.000 rpm).

### Temperature stage

The temperature stage is a custom-built system (Figure 2Α) and similar to the one used in previous studies.^76,77^ The setup is based on highly conductive sapphire glass (thermal conductivity of 27.1 W m^−1^ K^−1^, SMS-7521, UQG Optics) connected to two Peltier modules that are computer controlled via a proportional–integral–derivative (PID) device connected to the T-sensor. The sapphire quickly transmits the temperature of the Peltier to a transparent surface, enabling rapid active adjustment of temperature in the reaction chamber, which is pasted to the sapphire glass.

### TIRF assay, image acquisition and data analysis

TIRF microscopy assays were prepared as in ^19^. Briefly, coverslips used for microscopy were cleaned and silanized using dichlorodi-methylsilane, which renders the glass surface hydrophobic. Flow chambers were built using parafilms strips as spacers between a high thermal conductive sapphire slide and a silanized coverslip. All reactants flowed into the chamber were prepared in BRB80 buffer. First, neutravidin (100 mg/mL) was incubated for 5 min in the chamber to serve as linker. Then, the chamber was blocked using 1% w/v Pluronic F-127 (Sigma Aldrich) for 15 min. Washing occurred with 1 mg/mL K-casein. GMPCPP-stabilised seeds (10% Cy5-labelled and 20% biotin-labelled tubulin) were suspended in wash buffer (1 mg/mL K-casein) and flowed in the chamber with coverslip facing down. Reaction mix was prepared with 1 mg/mL k-casein, 1% β-mercaptoethanol, 2 mM Trolox, 2.5 mM PCA, 25 nM PCD, and 0.15% methyl-cellulose at different concentrations of purified tubulin with 10% Cy3- or Atto488-labelled porcine brain tubulin in BRB80. The chamber was sealed using two-component dental silicon (Picodent, Twinsil 22), and pasted to the temperature stage (sapphire against sapphire) using a thermally conductive paste (Keratherm KP 12). The reaction chamber was placed on the microscopy stage (coverslip facing the objective lens, temperature stage on top). A Nikon Ti-E microscope with a motorised TIRF angle was used, coupled to either an Andor DU-897 or a sCMOS, Photometrics Prime 95B camera.

Three TIRF datasets were acquired per condition for each species and were analysed manually by selecting each dynamic microtubule growing from the seed plus-end using Fiji.^78^ Kymographs were generated using a custom written macro involving the Multi Kymograph plugin (https://github.com/fiji/Multi_Kymograph). Quantitative parameters of dynamic instability were measured on the kymographs using the Fiji measurement tool. Catastrophe frequency was calculated using two methods. If shown as F_cat_ , it was calculated as the ratio of the total number of catastrophes within one assay, on the sum of all duration of growth events within that same assay. The values were averaged for each condition and plotted against tubulin concentration. A weighted linear regression was fitted to the data. If displayed as the average lifetime before catastrophe, they were calculated for each species by binning the growth events by velocity, and calculating the ratio of all the sum of growth events duration on the count of catastrophes for each bin.

Nucleation probability was calculated as the number of seeds that nucleated at least one microtubule within ten minutes of acquisition, divided by the total number of seeds within the field of view.

Plots were generated using the Altair package in Python.^79^ For all boxplots, the boxes range from 25 to 75th percentile, with whiskers spanning minimum to maximum values; the horizontal line marks the mean value. Linear regressions were performed using weighted least squares, the weights being the inverse of the value, unless stated otherwise.

### EB1 purification and comet length analysis

pETMM-11 vector containing His_6_-TEV-hEB1-GFP was expressed in *E. coli*. Expression was induced with IPTG, cells were grown overnight at 18°C. The next day, cells were pelleted, washed with 1xPBS and 5 mM EDTA, and resuspended in a lysis buffer. Upon addition of 0.5 mg/mL lyzozyme, cells were rocked at 4°C for 30 min and sonicated for 30 seconds, before being spun at 80.000 rpm in a MLA-80 rotor for 45 min at 4°C. The supernatant was filtered (0.22 µm) and flowed through an equilibrated Ni-sepharose column (HisTrap High Performance, Cytiva). The column was washed with a lysis buffer containing 10 mM imidazole and eluted with an elution buffer in 1 mL fractions. Buffer was exchanged to BRB80 containing 10% glycerol using a dialysis membrane, and concentration was determined by Nanodrop. 200 nM EB1-GFP were added to the TIRF-M assays as described above. All EB1-GFP assays were performed at 25°C. Tubulin concentration was adjusted depending on the species.

Comet lengths were analysed by first manually tracking comets within Fiji using the point tool. For each comet, its location at origin was selected on a movie slice, followed by the last occurrence of the comet on a later slice, both using the point tool of Fiji.^78^ The regions of interest created were exported to Python, together with the TIRF images sequence. A custom-written Python script, based on ^49^ was used to estimate comet length. Briefly, images were 6-time sub-sampled, a rectangle was cropped around the comet and rotated so that all comets were growing in the same direction. The intensity peak of each comet was aligned and assigned the coordinate 0, and for each comet the average growth velocity was calculated based on the total distance covered by the comet and the number of frames. Within each growth velocity, aligned pixels of each comet were averaged, the result was normalised, and all normalised average comets were super-averaged. The part of the profile starting one pixel (six subpixels) after the intensity peak and corresponding to the microtubule lattice was fitted to the following exponential decay function^80^:

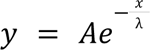

with A being the intensity at pixel a and λ the comet decay length. The peak pixel was excluded to avoid including potential subpixel perturbations within the tip structure. The rest of the profile was fitted to a Gaussian. The comet decay length λ was plotted against the average growth velocity calculated from each box.

### Intact protein analysis by liquid chromatography mass spectrometry and phospho-enrichment

Intact protein masses were determined by liquid chromatography mass spectrometry (LC-MS) as described in ^81^. In brief, samples were analysed using the Ultimate 3000 liquid chromatography system connected to a Q Exactive HF mass spectrometer via the ion max source with HESI-II probe (Thermo). The proteins were desalted and concentrated by injection on a reversed-phase cartridge (MSPac DS-10, 2.1×10 mm, Thermo). Full MS spectra were acquired using the following parameters: intact protein mode on, mass range m/z 600–2500, resolution 15,000, AGC target 3x10^6^, Microscans 5, maximum injection time 200 ms. The software tool UniDec was used for data processing and deconvolution of protein masses.^82^ First, an averaged spectrum was generated from each measurement followed by spectral deconvolution using the default settings except for the following adjustments: charge range 10-80, Mass range 48-52 kDa, sample mass every 1 Da, peak FWHM 0.1.

### Mass spectrometry peak assignment

Peaks from the mass spectrometry of tubulin samples from *X. laevis* and *X. tropicalis* were assigned as isoforms listed on Uniprot^83^ with the same mass (accession codes listed in Figure S6A-C). For *X. borealis*, although the genome is fully sequenced (assembly #GCA_024363595.1), there are no annotations that include genes, transcripts or tubulin protein sequences. We identified some coding regions for tubulin through NIH tBLASTn searches^84–86^ within the X. borealis genome (assembly #GCA_024363595.1) using *X. laevis* and *X. tropicalis* tubulin isoforms as references. Regions of the *X. borealis* genome surrounding these areas were exported (with ∼5,000 bases both upstream and downstream) and aligned within Benchling (https://benchling.com) to the equivalent areas for the corresponding tubulin isoforms within the *X. laevis* genome (Xenopus_laevis_v10.1 #GCF_017654675.1) and/or *X. tropicalis* genome (UCB_Xtro_10.0 #GCF_000004195.4). Splicing patterns within the *X. laevis* or *X. tropicalis* genomes were applied to identify and remove the orthologous intronic sequences from the *X. borealis* genomic sequence. The resulting synthetic mRNA transcripts were translated and used for mass spectrometry peak assignment and for structural modelling. Confidence in our approach was reinforced by the consistency of the predicted molecular weights to the mass spectrometry data, and the close similarity of the chromosomal locations and amino acid sequences of the isoforms between species.

### Cryo-EM sample preparation

Shortly before use, C-Flat 2/2 Holey Carbon grids (Protochips) were treated in air for 40 s at 0.2–0.3 Torr using a Harrick Plasma Cleaner. During the freezing process, grids were surface-treated in batches of two to minimise time between plasma cleaning and sample application. Grids were loaded into the humidity chamber of the Leica GP2 held at 95% humidity and at near-native temperatures for each sample (20/25/30°C for *Xl*, *Xb* and *Xt* respectively). Purified tubulin was quickly thawed and diluted with ice cold BRB80 to concentrations ranging from 8-16 µM of tubulin plus 1 mM GTP (see Table S1). 3 µL of the diluted tubulin was applied to the surface treated grid within the humidity chamber and this was incubated for 10 minutes to allow tubulin polymerisation. Excess liquid was removed from the grid through blotting with filter paper from the back of the grid (with the Leica GP2 sensor enabled distance of 0.3 mm and blotting time of 5 seconds) before immediate vitrification in liquid ethane. Blotting paper was changed ∼9 min into the incubation before blotting for each sample to minimise wrinkling from the humidity during the incubation.

### Cryo-EM grid screening

Grid screening images were collected on a Tecnai T12 transmission electron microscope (Thermo Fisher Scientific) operating at 120 kV with a US4000 4 K × 4 K CCD camera (Gatan). Images were recorded under low dose conditions at magnifications of 42,000 – 67,000 × corresponding to pixel sizes of 2.56 – 1.46 Å.

### Cryo-EM data collection

All high-resolution data were collected on a Titan Krios D3771 microscope (Thermo Fisher Scientific) operating at an accelerating voltage of 300 keV and using a BioQuantum K3 direct electron detector (Gatan) and post-column GIF energy filter (Gatan) with a slit width of 20 eV. The data were collected using the automated EPU software at a nominal magnification of 81,000 × with pixel size of 1.067 Å and a defocus range of -0.6 to -2.1 µm (Table S1).

### Cryo-EM data processing

Apart from particle picking, all image processing steps were carried out using RELION v3.1^87,88^ and the customised Microtubule RELION-based Pipeline (MiRP).^42,89^ Beam-induced motion within each movie was corrected using MotionCor2 within the RELION interface.^90^ The parameters of the contrast transfer function (CTF) for each motion-corrected, dose-weighted movie were determined using CTFFIND4 ^91^ again within the RELION interface. The resulting micrographs were imported into crYOLO v1.7.6 for automated filament picking.^92,93^ We generated an initial training model from coordinates of manually picked microtubules from a small subset of the corrected micrographs in the *X. borealis* dataset optimised to minimise picking of curved, flattened or overlapping microtubules. We used the same training model to pick data from all three samples with intervals of 81-82 Å between boxes, minimum filament lengths of 15-20 particles and thresholds optimised to each sample (Table S1). The resulting coordinates were imported into RELION with a box size of 544×544 pixels (580×580 Å) and 4x binning to 136×136 pixels.

4x binned particles were subjected to supervised 3D classification with low-pass filtered references of microtubules with different protofilament numbers. References were created using known helical parameters for different architectures.^74^ Using custom scripts,^42^ particles from each microtubule were then assigned a unified protofilament number class and separated by assigned architectures for downstream processing by a confidence threshold of 75% – filaments with confidence below this threshold were discarded at this stage. 14 and 15 protofilament microtubules with a 3-start helix were taken for further processing in each sample. In addition, helical 15 protofilament microtubules with a 4-start helix in the *X. laevis* dataset were also used for further processing.

### Microtubule architecture analysis

*2D width analysis:* All picked particles from each of the 3 species were extracted and run through a RELION 2D classification with 200 classes. Images of 120 most populated classes were exported for analysis using ImageJ^94^ following methods from ^95^. Briefly, using the plot-profile tool within ImageJ, we plotted the profile of a straight line drawn across the filament within each class average. The widths of each class average were determined by measuring the distance in pixels between the two peaks with highest intensity (indicating the edges of the MT). The widths recorded were then rounded to the nearest pixel, converted to Å (4.268 Å/px), and multiplied by the particle numbers within each class for representation in a histogram (Figure S3). The binning into the different protofilament numbers was defined using a published set of low-resolution EM reconstructions of bovine tubulin with a variety of different architectures.^74^

*3D class distribution:* As described above RELION 3D classification was used to separate particles according to their helical architectures using suitable references. *TubuleJ* ^38^: This analysis was done on individual microtubule images following low-dose imaging during the grid screening process (see section above). Individual microtubules with the clearest lattices were selected by eye for helical lattice parameter analysis in TubuleJ.^38^

### 3D reconstruction

The particles, separated by helical architecture, were re-extracted with 2x binned and a smaller box size of 208x208 pixels (444×444 Å) for downstream processing (Figure S4 B). Following previously published MiRP processing pipelines,^42^ we performed several rounds of 3D alignment followed by smoothing of Euler angles and X/Y shift assignments based on the prior knowledge of MT architecture using Python(v2.7.5) scripts were performed. From this, an initial seam location was determined for each MT, which was then checked and corrected using supervised 3D classification against references for all possible seam locations. For all three species, particles assigned into 14-3 and 15-3 classes at the protofilament sorting steps, were treated similarly (Figure S4 C-F). The particles classified as fully helical (15-4 microtubules from *X. laevis*) were treated slightly differently as seam alignment and correction steps were skipped, while the absence of a seam subsequently allowed imposition of helical symmetry during refinement.

Following the MiRP alignment steps, 2D classification was performed for all architectures (with no image alignments) and any overlapping microtubules picked by crYOLO were discarded. Particles were re-extracted without any binning for RELION refinement (1.067 Å/px) without symmetry impositions (C1 symmetry) and restricted Euler searches (Figure S4C). Aligned particles were further processed through iterations of RELION Bayesian Polishing and CTF refinement,^96,97^ focussed 3D classification and (for 3-start microtubules) symmetry expansion by relion_particle_symmetry_expand (Figure S4D).^98,99^ Focussed 3D classification without alignments (with masking for a single protofilament opposite the seam) allowed correction of misaligned α- β-tubulin register and removal of poorly aligned pseudo-particles. RELION 3D auto-refinements were then repeated for a full cross-section of the microtubule (Figure S4E) and with a mask around only 6 tubulin dimers opposite the seam (Figure S4F). RELION post-processing of unsharpened half-maps from each refinement was used to generate Fourier Shell Correlation (FSC) analysis to describe resolution (Figure S4 E,F). Finally, we ran unsharpened half-maps from each refinement through deepEMhancer for local sharpening to improve the interpretability for model building (Figure S5 A).^75^

### Model generation, refinement and analyses

Initially AlphaFold2 multimer was used to generate the model for a single tubulin dimer from *X. tropicalis* with the amino acid sequences for the tubulin isoforms most represented within mass spectrometry (TUBAL3.2:TUB4B).^100–102^ The AlphaFold model, GDP, GTP and magnesium atoms were rigid-body docked into their corresponding densities for the 14-3 microtubule from *X. tropicalis* using Fit-in-Map in ChimeraX.^103,104^ Initial model refinement for a single tubulin dimer was conducted using interactive molecular dynamics force field-based model fitting in ISOLDE within ChimeraX,^105^ against both the deepEMhancer post-processed density (Figure S5A) and unsharpened reconstructions from RELION 3D auto-refine (Figure 4F). Finally, a model for 6 tubulin dimers was fit to the density and refined through iterations of Phenix real-space refinement with non-crystallographic symmetry restraints and validation and correction within ISOLDE. A single dimer from the refined *Xt* model was then used as a starting model for 14-3 microtubules in *Xb* and *Xl* by altering the amino acid sequence using the swapaa tool in ChimeraX. Modelling was repeated as described above. Finally, the 14-3 models were used as the starting models for 15-3 and 15-4 architectures within the same species and refined using Phenix real-space refinement as above.

### Model analysis – 6 dimer models

The RMSD depictions between models (Figure 5 A,E,F) were calculated using ChimeraX following alignment of models on the N-terminal domain (residues 1-207) of the indicated β,-tubulin. Dimer rise, inter- and intra-dimer spacing were determined by distance measurements between centroids at the centre of mass for each tubulin monomer within ChimeraX. The CA-CA distance measurements (Figure 5 K-L) between residues were determined from pdb files for each model using custom python scripting generated using ChatGPT.^106^ The interaction analysis for the resulting models (Figure B-D, G-J) was conducted using the ’protein interfaces, surfaces and assemblies’ (PISA) service at the European Bioinformatics Institute (https://www.ebi.ac.uk/pdbe/pisa/).^45^

### Complete lattice comparisons

To compare models for 14-3 microtubule lattices across datasets (Movie S1) we selected a subset of ∼15,000 particles that had 70% confidence following seam alignment steps in the MiRP pipeline (see above).^42^ Selected particles were aligned through RELION auto-refine either without any symmetry impositions (C1 symmetry), or with full helical symmetry imposed. Complete microtubule lattice models were assembled through rigid-body docking of individual tubulin monomers into the symmetrized and non-symmetric reconstructions. The two models were then aligned and root-mean square deviations between them were coloured within ChimeraX.^103,104^ Images were exported from ChimeraX and movies were generated in Microsoft PowerPoint.

### Arrhenius equation

According to a simple microtubule growth model *T* + *T_n_* ⇌ *T_n+1_*, the linear relationship between growth velocity and tubulin concentration follows the equation: v_g_ = k_on_[T] − k_off_ with v_g_ being the growth velocity, k_on_ the on-rate constant and k_off_ the off rate. Using the k_on_ for temperature and species, we plotted the linear Arrhenius equation: ln(k_on_) = ln(A) − E_a_.RT.

## QUANTIFICATION AND STATISTICAL ANALYSIS

In each figure legend, details about the quantifications have been provided, including the number of events measured (n), the mean/median values, the SD/SEM. In addition, information about the statistical tests used for measuring significance and interpretation of p values is provided. P values greater than 0.05 are represented by “ns”. A single * indicates a p value ≤ 0.05, ** indicates p values ≤ 0.01, *** indicates p values ≤ 0.001, and **** indicates p values ≤ 0.0001. For statistical analysis and plotting in this paper, we utilised Graphpad Prism version 8.0 for Mac OS X, GraphPad Software, La Jolla California USA, https://www.graphpad.com. When necessary, graph visuals such as line thickness, fonts, and colours were optimised using Adobe Illustrator. Data visualisation was performed in the Python Altair library.^79^

## RESOURCE AVAILABILITY

### LEAD CONTACT

Further information and requests for resources and reagents should be directed to and will be fulfilled by the lead contact, Simone Reber.

### MATERIALS AVAILABILITY

This study generated purified *X. borealis* tubulin as a unique reagent.

### DATA AND CODE AVAILABILITY

All data needed to evaluate the conclusion in the paper are present in the paper and the supplementary materials. The code CometTrack to analyse the EB1-GFP comet length is available on github: https://github.com/elladegaulejac/CometTrack. Depositions of structural data are summarised in Table S1: Cryo-EM reconstructions for 14-3, 15-3, and 15-4 microtubules from *X. laevis*, *X. borealis*, and *X. tropicalis* have been deposited in the Electron Microscopy Data Bank under accession codes EMD-50811, EMD-50941, EMD-50942, EMD-50943, EMD-50945, EMD-50946, and EMD-50947; corresponding atomic models for 6 tubulin dimers have been deposited in the Worldwide Protein Data Bank under accession codes 9FVJ, 9G0O, 9G0P, 9G0Q, 9G0R, 9G0S, and 9G0T.

## ACKNOWLEDGEMENTS

We thank former and current members of the Moores and Reber laboratories for critical discussion, in particular William Hirst, Soma Zsoter, and Harry York who helped with initial data collection. We thank Gil Henkin for critical comments on the manuscript and Juan Iglesias-Artola and Anatol Fritsch for building the sapphire stage supported by MAX!mize, the official start-up incubation program for the Max Planck Society. We thank the AMBIO (Charité, Berlin) for imaging support. For mass spectrometry, we would like to acknowledge the assistance of the Core Facility BioSupraMol supported by the Deutsche Forschungsgemeinschaft (DFG). This work was supported by the Max Planck Society (S.R.). This article was prompted by a stay at the Marine Biological Laboratory (MBL), Woods Hole, MA funded by the Princeton-Humboldt Strategic Partnership Grant. Cryo-EM work was supported by the Medical Research Council, U.K. (MR/R00352/1 and MR/Y000633/1 to C.A.M.). Cryo-EM data were collected at ISMB EM facility (Birkbeck College, University of London) with financial support from the Wellcome Trust (202679/Z/16/Z and 206166/Z/17/Z). J.S. is supported by a Bloomsbury Colleges PhD Studentship (LIDo program). We thank N. Lukoyanova and S. Chen for electron microscope support and D. Houldershaw for computational support at Birkbeck. L.T., J.S. and C.M. acknowledge I. Nobeli’s support and useful discussions concerning sequence analysis.

## AUTHOR CONTRIBUTIONS

L.T. performed all structural experiments including model building. E.G. purified all tubulins, performed and analysed all *in vitro* experiments. A.B. provided egg extracts for tubulin purification. B.K. performed and interpreted the mass-spectrometric analyses, peak assignment by E.G. J.S. worked with L.T. on *X. borealis* sequence analysis. C.M. supervised structural data collection and analysis and gave critical inputs throughout the project. S.R. conceptualised the study. Visualisation by L.T., E. G., and S.R. L.T. and S.R. wrote the final manuscript with input from all authors.

## DECLARATION OF INTERESTS

The authors declare no competing interests.

## Notes

### Competing Interest Statement

The authors have declared no competing interest.

